# Aberrant medial entorhinal cortex dynamics link tau pathology to spatial memory impairment

**DOI:** 10.64898/2025.12.30.696887

**Authors:** Taylor J. Malone, Kyle Cekada, Jean Tyan, Lujia Chen, Garret Wang, Yi Gu

## Abstract

Tau pathology in the entorhinal cortex (EC) is associated with spatial memory decline in aging and early-stage Alzheimer’s disease, but its impact on EC computations during learning is not well understood. We performed longitudinal two-photon calcium imaging of layer 2 excitatory neurons in the medial EC (MEC) of PS19 tauopathy mice over 10 days of an operant spatial learning task. Male PS19 mice showed marked learning impairments accompanied by dysregulated MEC activity and unstable spatial coding. Their activity also showed weakened representations in cue-poor relative to cue-rich regions, correlated with attenuated speed modulation. These changes suggest that impaired path integration destabilizes MEC spatial maps, leading to impaired spatial memory. In contrast, female PS19 mice exhibited only mild behavioral and neural deficits despite a comparable tau burden, suggesting sex-specific resilience. Among MEC cell types, pyramidal cells accumulated more phosphorylated tau than stellate cells and displayed the most severe functional disruption, linking cellular tau load to circuit dysfunction. Finally, general linear models of MEC activity reliably predicted learning performance, highlighted particularly strong contributions from non-grid and pyramidal cells, and accurately classified PS19 versus wild-type mice. These findings identify aberrant MEC dynamics as a key circuit mechanism underlying tau-related spatial memory deficits and point to early diagnostic and circuit-targeted therapeutic strategies.

## INTRODUCTION

Tau pathology, characterized by the aggregation of microtubule-associated protein tau (MAPT), is a defining feature of Alzheimer’s disease (AD) and other tauopathies^1,2^. In the earliest stages of AD and aging, tau pathology prominently involves medial temporal lobe structures, particularly the entorhinal cortex (EC), which is among the earliest cortical regions to show neurofibrillary tangles (NFTs) and hyperphosphorylated tau inclusions^3–5^. The EC is a central hub within the hippocampal-entorhinal circuit^6^ and is crucial for spatial representation and memory, as demonstrated by fMRI and lesion studies in humans and rodents^7–13^. Consistent with this role, early AD is characterized by deficits in encoding^14–18^, consolidating^19^, and retrieving^16–21^ spatial memories, as well as in allocentric^14,15,22–25^ and egocentric^14,22,24,25^ spatial representations that support cognitive map construction. In addition, NFT accumulation increases with age independently of AD status, becoming highly prevalent in older individuals, and correlates with spatial memory impairment during normal aging^26–28^. In contrast, amyloid-β, another hallmark of AD, shows less regional specificity for the EC in early AD and is less prevalent in the EC than tau pathology in normal aging, with many older individuals showing substantial entorhinal tau in the relative absence of amyloid-β^4,5,26,29,30^. Amyloid-β pathology is also less predictive of memory deficits^27,31^. Collectively, these findings implicate the EC as an early locus linking tau pathology to spatial memory decline in normal aging and AD.

Additional support for this hypothesis comes from studies on EC neural dynamics. In rodents, the medial entorhinal cortex (MEC) contains a diverse population of functional cell types that encode spatial and self-motion information^32–38^. Among these, grid cells are the most extensively studied and fire in a triangular lattice pattern in open arenas^32^. AD mouse models that recapitulate tau pathology, with or without amyloid-β, reveal widespread disruption of MEC activity^39–42^. Grid cells in these models exhibit hypoactivity, disrupted field periodicity, reduced spatial information, and impaired speed modulation^39,40^. The activity of non-grid cells and interneurons is also affected to various extents^39,40^. Additionally, *in vitro* electrophysiology studies report altered intrinsic firing properties of MEC cells^41,42^ with variability across age^42^, subregion^41^, and cell type^42^. Lastly, human fMRI studies reported reduced grid-cell-like activity in the EC of older adults and of young adults with genetic risk for AD^9,11^.

Despite these advances, it remains unclear how EC neural activity contributes to spatial memory deficits associated with tau pathology. Current studies suggest that spatial memory formation comprises a rapid initial encoding phase followed by several days of memory consolidation^43–46^, during which behavioral performance gradually improves^47–49^. To date, however, no studies have longitudinally tracked MEC activity across the full course of this process and related MEC activity to spatial learning performance in the context of tau pathology. Instead, most prior work has examined MEC activity at isolated navigational time points^39–42^. Furthermore, some studies have investigated MEC activity under conditions of high amyloid-β burden^50–54^ or severe neuronal loss^39^. The presence of amyloid-β can obscure tau-specific effects on neural activity, particularly in aging-related tau pathology. Severe neuronal loss is also less representative of aging and early AD, when overt atrophy is not yet present as in symptomatic AD and other tauopathies^55–57^, and may confound the specific effects of tau pathology on neuronal function at the cellular level through network-wide disruption. Together, these limitations hinder the identification of MEC neural activity features that contribute to deficits across the full trajectory of memory formation under tau pathology in aging and early AD.

To address this gap, we used two-photon imaging to longitudinally track calcium dynamics of layer 2 MEC neurons in the PS19 tauopathy mice^58^ during an operant spatial learning task across 10 days in virtual reality (VR) environments. PS19 mice express human tau bearing the P301S mutation that causes familial frontotemporal dementia in humans^1,59^ and are widely used to model tau pathology relevant to AD^60^, as they develop robust, progressive tau accumulation from ∼5–6 months of age across multiple brain regions^58,61^, including the MEC^61^, followed by deficits in hippocampal- and entorhinal-dependent spatial memory at 6-8 months^60–65^. More pronounced hippocampal–entorhinal atrophy typically emerges later, around 9–12 months^58^, allowing us to examine how MEC neural dynamics contribute to spatial deficits before prominent neuronal loss, a disease stage that more closely resembles aging and early AD.

We found that compared to wild-type (WT) mice, male PS19 mice demonstrated severely impaired spatial learning. Their MEC activity showed reduced speed modulation, lower activity map stability within and between days, and failure to develop a spatial map shared by other groups. Their MEC activity was abnormally anchored to visual cues, suggesting disrupted path integration. In contrast, female PS19 mice showed limited spatial learning and MEC activity deficits. Activity deficits were strongest in pyramidal cells, which accumulated more phosphorylated tau than stellate cells, particularly in males. Finally, generalized linear modeling linked altered MEC activity—especially in non-grid and pyramidal cells—to spatial learning performance and reliably distinguished PS19 from WT males. Together, these findings highlight EC circuit dysfunction as a key contributor to tau-pathology-related spatial memory deficits in aging and early AD and identify the EC as a potential target for early diagnostic and therapeutic interventions.

## RESULTS

### Pyramidal cells in PS19 mice show increased tau pathology

To allow long-term monitoring of calcium activity during spatial learning, we crossed PS19 mice^58^ with Thy1-GCaMP6f transgenic mice (GP5.3) to drive stable expression of the calcium indicator GCaMP6f in excitatory neurons in MEC layer 2^66–68^, a region particularly vulnerable in AD^69,70^. F1 offspring expressing both P301S tau and GCaMP6f were designated as “PS19 mice” and littermates expressing GCaMP6f but not P301S tau were WT controls. Young PS19 (up to 6 months old) and WT mice displayed low immunoreactivity to the AT8 antibody (**Figs. 1A-C**), which recognizes tau phosphorylated at serine 202 and threonine 205^71^ (pTau), an early pathological tau species that is strongly enriched in NFTs^72^. In contrast, immunoreactivity strongly increased in older PS19 mice (> 6 months) (**Figs. 1A-C**), consistent with reported NFT emergence around 5 months of age^58^. Given the well-documented sex differences in patients with AD and tauopathy models^73–77^, we examined whether overall pTau accumulation varied between male and female mice. Although no significant differences were observed, females exhibited a trend toward broader pTau spread and but lower local pTau intensity (**Figs. 1D, E**).

**Figure 1:**
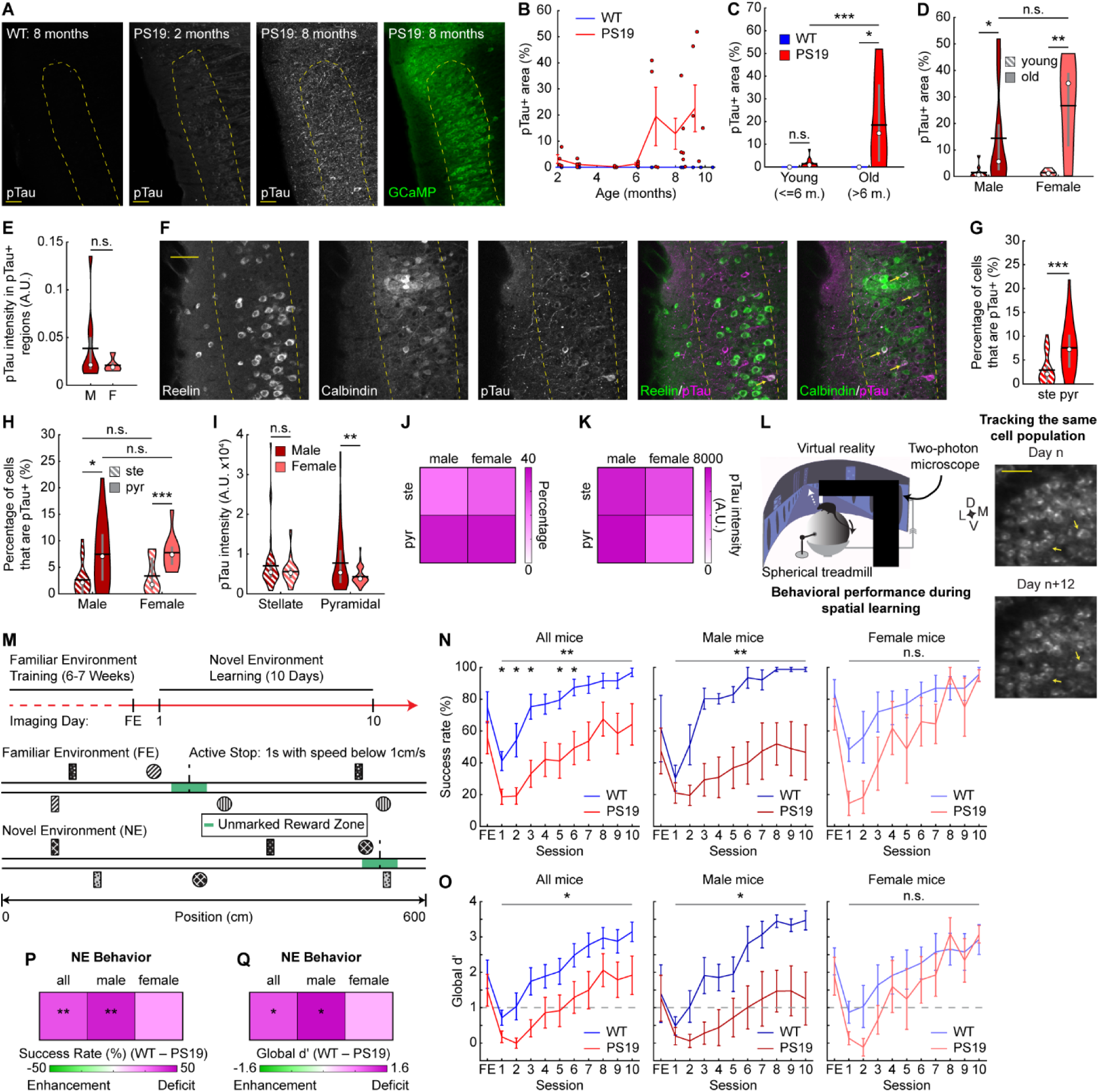
Male PS19 mice show behavioral deficits in an operant spatial learning task. **A.** Left: Example images showing pTau accumulation in WT and PS19 mice of different ages. Right: GCaMP6f in an 8-month PS19 mouse. Dotted yellow line indicates MEC layer 2. Scale bar: 50µm **B.** Quantification of pTau accumulation in MEC layer 2 as a function of age. Dots represent the exact age of individual mice. Lines represent ages pooled into similarly aged clusters. **C.** pTau accumulation in MEC layer 2 in young (age<=6 months, m.) and old mice (age>6 months). Unpaired Student’s t-test. **D.** pTau accumulation in MEC layer 2 in young and old PS19 mice split by sex. Unpaired Student’s t-test. **E.** Average pTau intensity in pTau positive regions of MEC layer 2 in old male (M) and female (F) PS19 mice. Unpaired Student’s t-test. **F.** Example images showing the overlap of reelin (stellate cell marker) and calbindin (pyramidal cell marker) with pTau accumulation. Dotted yellow line indicates MEC layer 2 boundaries. Arrows indicate overlap with pTau. Scale bar: 50µm **G, H.** Percentage of stellate (ste) and pyramidal (pyr) cells overlapping with pTau+ cells in all PS19 mice (**G**) or in PS19 mice split by sex (**H**). Paired/Unpaired Student’s t-test. **I.** pTau intensity in stellate and pyramidal cells that overlap with pTau+ cells in PS19 mice split by sex. **J.** Heatmap summarizing stellate and pyramidal cells overlapping with pTau+ cells in PS19 mice by sex and cell type. Color represents mean value in **H**. **K.** Heatmap summarizing pTau intensity in stellate and pyramidal cells overlapping with pTau+ cells in PS19 mice. Color represents mean value in **I**. **L.** Experiment schematic. Left: virtual reality system, adapted from^68^. Right: Example field-of-view (FOV) of calcium imaging showing reliable cell tracking across days. Arrows point to examples of tracked cells. D: dorsal; V: ventral; M: Mediolateral; L: Lateral. Scale bar: 50µm. Full FOV is 750µm x 750µm. **M.** Task schematic. **N.** Percentage of laps that mice successfully stopped to receive water reward across learning in all mice (left), male mice (middle), and female mice (right). **O.** Global reward zone discrimination (global d’) across learning in all mice (left), male mice (middle), and female mice (right). Horizontal dotted line represents learning threshold. **N, O.** Horizontal gray bars indicate p values for the group difference (two-way repeated measures ANOVA). Individual time points are compared using Student’s t-test with Bonferroni-Holm correction. **P, Q.** Heatmaps summarizing the success rate (**P**) or global d’ (**Q**) difference between WT and PS19 mice by sex. Color represents the mean difference between WT and PS19 values averaged across days 1-10 in **N** and **O**, respectively. Unpaired Student’s t-test. \**p* ≤ 0.05, \*\**p* ≤ 0.01, \*\*\**p* ≤ 0.001. In violin plots, horizontal line represents mean, white circle represents median, and whiskers represent interquartile range. Error bars in line plots represent mean ± sem. Statistical details can be found in **Table S1**. See also **Figs. S1, S2**.

We next examined pTau accumulation in stellate and pyramidal cells, which are the primary excitatory cell types in MEC layer 2 and differ in their circuit connectivity, contributions to spatial navigation, and susceptibility in AD animal models and human patients^42,78–84^. Quantifying the overlap of pTau with reelin (a stellate cell marker) and calbindin (a pyramidal cell marker), we found that pyramidal cells were more often pTau+ than stellate cells (**Figs. 1F, G**), and pTau+ cells were more likely to be pyramidal than stellate cells (**Fig. S1A**). This pyramidal enrichment was evident in both sexes (**Figs. 1H, S1B-D**), but pTau intensity within the overlapping pyramidal cells was higher in males than females (**Fig. 1I**), consistent with higher local pTau abundance in males (**Fig.1E**). These outcomes were summarized in 2-by-2 heatmaps by cell type and sex (**Figs. 1J, K**). Overall, these results suggest that pyramidal cells are a particular target of tau pathology in MEC layer 2 of PS19 mice, especially in males.

To investigate MEC neural dynamics underlying spatial memory of PS19 mice, we used mice at ages showing spatial memory deficits and pTau accumulation but preceding the hindlimb paralysis and neurodegeneration reported in this line^58,63^. Hindlimb paralysis could impair running behavior during navigation, whereas neurodegeneration could confound cell-autonomous effects of pTau on neural function by disrupting network connectivity. Previous studies indicate that PS19 mice develop spatial memory deficits at approximately 6-8 months of age, when mobility remains largely intact^60,62–65^. In our F1 mice, significant pTau accumulation began after 6 months (**Figs. 1B-D**) and the median age of paralysis onset was 11.6 months (**Fig. S1E**). Accordingly, all experiments were conducted in mice aged 7–10 months, prior to paralysis onset (**Fig. S1F**). Quantification of total number of stellate and pyramidal cells per area revealed no significant neurodegeneration at these ages overall (**Fig. S1G)** or by sex (**Fig. S1H**).

Thus, all subsequent experiments used 7-10-month-old mice. At these ages, PS19 mice show robust pTau accumulation, particularly in layer 2 pyramidal cells, without significant paralysis or neuronal loss. The robust colocalization between pyramidal cells and pTau has also been observed in human brains in early stages of AD^83^, prompting us to investigate the function of these neurons in contrast to stellate cells.

### Male PS19 mice show behavioral deficits in an operant spatial learning task

To directly investigate the effects of tauopathy on MEC activity during spatial learning, we performed cellular-resolution two-photon calcium imaging in layer 2 of the MEC in 21 head-fixed mice (10 WT, 11 PS19) during VR navigation (**Fig. 1L**), enabling longitudinal tracking of hundreds of neurons per genotype group across multiple days of learning (**Fig. 1L**). Water-restricted mice were trained to navigate a 6-meter familiar environment (FE) and stop in an unmarked 50-cm reward zone to trigger a water reward delivery (**Fig. 1M**). After learning the task, mice were imaged for one day in the FE and for 10 consecutive days while learning to navigate a novel 6-meter environment (NE) with new visual cues and a water reward at distinct locations compared to the FE (**Fig. 1M**). During the FE training, PS19 mice experienced a similar number of training days (**Fig. S1I**) and traversed more laps than WT mice (**Fig. S1J**), demonstrating at least as much opportunity to learn the task. To ensure comparable NE experience, NE sessions were limited to 20 laps per day (**Figs. S1I, J**).

Learning was quantified by success rate, the percentage of laps rewarded. PS19 mice exhibited significantly lower success rates than WT mice during spatial learning (**Figs. 1N, P**). When analyzed by sex, PS19 males showed marked learning deficits, whereas PS19 females showed a mild, nonsignificant trend toward impairment (**Figs. 1N, P**). WT mice also learned faster (learning rate constant, WT*_K_* or PS19*_K_*) and reached 95% of plateau earlier (WT_95%_ or PS19_95%_) than PS19 mice (**Fig. S2A**). To test whether mice were obtaining reward through indiscriminate stopping, we quantified the discrimination between the reward zone and other track locations using global d’^85,86^, with global d’>1 indicating successful discrimination^85–87^. Both WT and PS19 mice eventually exceeded this threshold, but WT mice displayed higher overall discrimination (**Figs. 1O, Q**) and reached the threshold sooner (**Fig. S2B**). The discrimination deficit was significant in PS19 males but not females (**Figs. 1O, Q**), while both sexes showed a trend of delayed rise to the threshold (**Fig. S2B**). Success rate and global d’ were also highly correlated (**Fig. S2C**), further arguing against an indiscriminate stopping strategy.

Because hyperlocomotion (i.e., elevated locomotor activity) has been reported in PS19 and other tauopathy and amyloid-β mouse models^60,88–90^, we tested whether impaired stopping could explain their spatial learning deficits. PS19 mice made similar numbers of stops as WT mice (**Fig. S2D**), indicating intact stopping, and still showed fewer stop attempts in the reward zone even under lenient stopping criteria (expanded reward zone, increased speed threshold, and shortened time to define a reward-zone stop) (**Fig. S2E**). Finally, while PS19 mice displayed a nonsignificant trend toward faster movement (**Fig. S2F**), when a subset of WT and PS19 mice were matched by their velocity on a per-day basis (**Fig. S2G**), PS19 mice still exhibited significantly lower success rate and global d’ than WT mice, especially among males (**Figs. S2H, I**).

In summary, PS19 mice exhibit significant spatial learning impairments, characterized by lower success rates and weaker reward-zone discrimination. This deficit was particularly pronounced in male mice, while PS19 females showed only a non-significant trend toward impairment but reached peak success more slowly than WT females. These learning impairments were not explained by stopping deficits or hyperlocomotion, suggesting that disrupted neuronal activity likely underlies the spatial learning deficits in PS19 mice.

### MEC cells of PS19 mice show hyper- or hypoactivity depending on molecular identity and sex

We first compared the molecular identity (stellate vs. pyramidal) of active cells in MEC layer 2 of PS19 and WT mice. For the FE day, GCaMP6f+ stellate and pyramidal cells were identified by soma size inferred from GCaMP6f fluorescence^68,91^ (**Figs. S3A-D**). Compared to WT mice, PS19 mice had more active cells per square millimeter (**Fig. 2A**), a trend toward a higher fraction of active cells—particularly stellate cells— (**Fig. 2B**), and a lower pyramidal-to-stellate ratio among active cells (**Fig. 2C**). These trends were similar across sexes (**Figs. S3E-H**). Thus, PS19 mice showed a trend toward MEC hyperactivity driven by increased active cell numbers and stellate cell enrichment.

**Figure 2:**
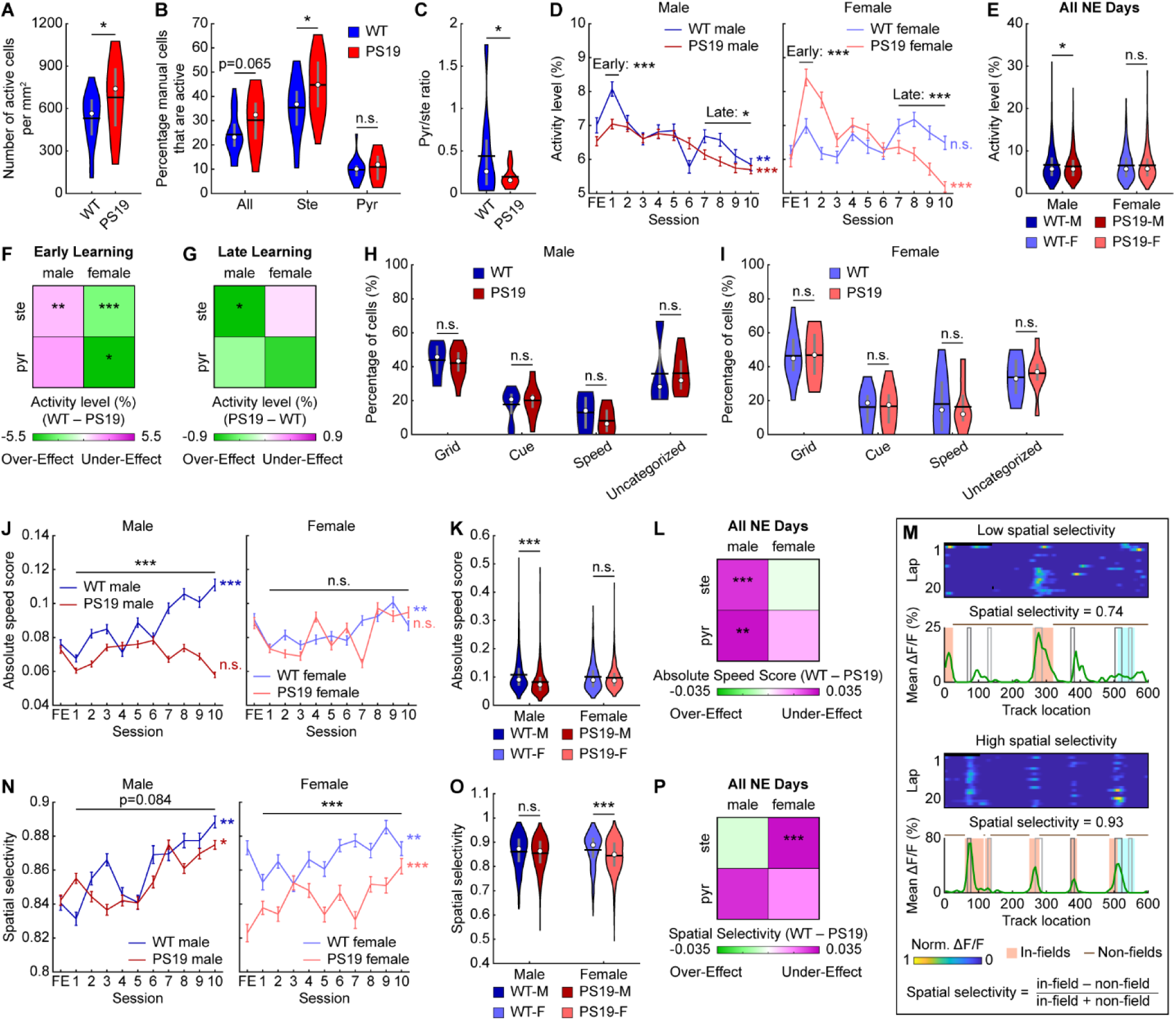
The MEC of PS19 mice exhibit altered neural activity level, speed modulation, and spatial selectivity. **A.** The number of active cells per area on the FE day. **B.** The percentage of all manually outlined cells and manually outlined stellate (ste) and pyramidal (pyr) cells that are active on the FE day. **C.** The ratio between the fraction of active cells that are identified as pyramidal cells and as stellate cells. A low value indicates stellate cell enrichment. **D.** Activity level across learning in male (left) and female (right) mice. **E.** Activity level averaged across all NE days. WT-M: WT Male, PS19-M: PS19 Male, WT-F: WT Female, PS19-F: PS19 Female **F, G.** Heatmaps summarizing the activity level difference between WT and PS19 mice by sex and cell type. Color represents the mean difference between WT and PS19 values averaged across early (day 1, **F**) and late (days 7-10, **G**) learning. Unpaired Student’s t-test. **H, I.** The percentage of common cells identified as grid, cue, speed, or unclassified cells in male (**H**) and female (**I**) mice. **J.** Absolute speed score across learning in male (left) and female (right) mice. **K.** Absolute speed score averaged across all NE days. **L.** Heatmap summarizing the absolute speed score difference between WT and PS19 mice by sex and cell type. Color represents the mean difference between WT and PS19 values averaged across all learning days as in **Fig. S4D**. Unpaired Student’s t-test. **M.** Schematic of spatial selectivity calculation. Top: Calcium activity of example cells as a function of lap and track location with low (top) and high (bottom) spatial selectivity. Bottom: spatial selectivity calculation for the above cells based on activity level across laps as a function of track location. Black and grey rectangles represent left and right cues, respectively. Blue rectangle represents reward zone. **N.** Spatial selectivity across learning in male (left) and female (right) mice. **O.** Spatial selectivity averaged across all NE days. **P.** Heatmap summarizing the spatial selectivity difference between WT and PS19 mice by sex and cell type. Color represents the mean difference between WT and PS19 values averaged across all learning days as in **Fig. S4H**. Unpaired Student’s t-test. \**p* ≤ 0.05, \*\**p* ≤ 0.01, \*\*\**p* ≤ 0.001. In violin plots, horizontal line represents mean, white circle represents median, and whiskers represent interquartile range. Violin plots use unpaired Student’s t-test. Error bars in line plots represent mean ± sem. In line plots, horizontal gray bars indicate p values for the group difference (early learning: unpaired Student’s t-test, late learning: two-way repeated measures ANOVA, all learning days: linear mixed-effects model), and p-values to the right indicate significant Pearson correlation of the mean value with respect to time. Statistical details can be found in **Table S1**. See also **Figs. S3, S4**.

We further examined whether single-cell activity levels differed by calculating the daily average temporal activity (mean ΔF/F) in common cells (neurons active across all 11 imaging days: 1 FE day and 10 NE days). Because common cell activity is strongly associated with spatial learning^49^, these cells likely contribute to spatial memory formation. In most groups, activity increased sharply on NE day 1 and then gradually declined across learning days. In contrast, PS19 males showed only a modest increase on day 1 and lower activity than WT males on subsequent days, resulting in an overall reduced activity level (**Figs. 2D, E**). PS19 females, however, exhibited a larger activity increase in early learning (day 1) followed by lower activity on late learning days (days 7-10) (**Fig. 2D**), yielding comparable overall activity to WT females (**Fig. 2E**). These patterns were largely consistent across stellate and pyramidal cells, although cells in PS19 females showed mixed trends relative to WT during later learning (**Figs. S3I, J)**. Stellate cells in male and female PS19 mice showed strong trends of overall hypoactivity and hyperactivity, respectively, with weaker trends in pyramidal cells **(Fig. S3K**).

To relate activity differences to the cell-type- and sex-specific tau accumulation pattern (**Figs. 1J, K**), we summarized molecular cell-type- and sex-specific activity level differences between PS19 and WT mice using 2×2 heatmaps for early and late learning (**Figs. 2F, G**). Green and magenta indicate over- and under-effects in PS19 mice, defined as activity exceeding or failing to reach the expected WT activity trend, respectively. Because WT activity increased on day 1 (**Figs. 2D, S3I, J**)^49^, lower and higher PS19 activity on day 1 relative to WT mice indicated under-and over-effects, respectively (**Fig. 2F**). During late learning, when WT activity declined (**Figs. 2D, S3I, J**), lower and higher PS19 activity indicates over- and under-effects, respectively (**Fig. 2G**). Importantly, both over- and under-effects could lead to cognitive deficits, as they indicate activity deviations from the WT level. Subsequent 2x2 heatmaps were generated using the same definitions.

In summary, PS19 mice tend to have more active cells in the MEC, with stellate cell enrichment relative to WT mice. Given the greater tau accumulation in pyramidal cells, stellate cell enrichment may reflect compensation for pyramidal dysfunction. However, mean activity levels differ from WT in a cell-type-, sex-, and learning-stage-dependent manner.

### MEC neurons of PS19 mice have altered speed modulation and spatial selectivity

We further investigated functional cell type composition of common cells and their activity changes with respect to speed and spatial location, key regulators of MEC neural activity^32,34,37,38^. We classified grid cells, neurons active in a triangular lattice pattern in a two-dimensional (2D) space^32^, during the navigation of our one-dimensional (1D) VR tracks^68,92,93^. We also classified cue cells, which fire around visual landmarks^36^, and speed cells, whose activity is significantly correlated with mouse velocity^37,38^. The common cell population of each functional cell type was those cells that met the classification criteria on ≥6 imaging days (more than half of the days)^49^. The remaining cells were considered unclassified. Functional cell type proportions were largely similar across genotypes in both sexes (**Figs. 2H, I**). The most notable difference was a trend toward fewer speed cells in PS19 males (**Figs. 2H**), suggesting reduced speed modulation.

To directly measure speed modulation, we calculated speed scores, the correlation between single-cell temporal activity and velocity^37,38^. In WT males, raw speed scores remained relatively stable across learning (**Fig. S4A**) because both positive and negative speed scores gradually increased in magnitude (**Figs. S4B, C**). This pattern was absent in PS19 males, which showed no magnitude increases (**Figs. S4B, C**) and lower raw and signed speed scores **(Figs. S4A-C)**, particularly for positively modulated cells. In contrast, WT females showed decreasing negative speed scores (i.e., more negatively modulated activity by speed), leading to decreasing raw speed scores (**Figs. S4A-C**). PS19 females followed the same trends (**Figs. S4A-C**). Absolute speed scores, which reflect modulation strength independent of direction, tended to increase across learning in WT mice of both sexes and in PS19 females, but not in PS19 males (**Fig. 2J**). PS19 males also exhibited significantly lower absolute scores than WT males (**Figs. 2J, K**), including in stellate and pyramidal cells (**Figs. 2L, S4D**). To assess functional cell-type differences, we analyzed common grid cells versus non-grid cells (all other common cells) and found lower absolute speed scores in PS19 males than WT males in both cell types (**Fig. S4E**). Because absolute speed scores rise during learning in WT mice (**Fig. 2J**), reduced speed modulation in PS19 males represents an activity deficit.

Furthermore, we assessed spatial selectivity because many cells exhibited spatial fields, clusters of adjacent spatial bins with activity above chance (**Fig. 2M**). Spatial selectivity was quantified by in-field versus non-field activity (**Fig. 2M**). Male PS19 mice exhibited similar selectivity to WT mice, whereas PS19 females showed lower selectivity (**Figs. 2N, O**), primarily due to elevated non-field activity that outweighed their slightly higher in-field activity (**Figs. S4F, G**). This reduction was largely attributable to stellate rather than pyramidal cells (**Figs. 2P, S4H**) and was present in both grid and non-grid cells (**Fig. S4I**). Because spatial selectivity increased with learning (**Fig. 2N**), lower spatial selectivity in PS19 females likely represents an activity deficit.

Together, these results revealed activity differences between PS19 and WT mice that vary by molecular and functional cell types. The diminished speed encoding in males and the reduced spatial selectivity in female PS19 mice may underlie their impairments in spatial learning.

### Male PS19 mice have impaired MEC within-day and cross-day activity consistency

Spatially consistent MEC activity is essential for successful spatial learning^49^. To assess whether PS19 mice exhibit deficits in spatial activity consistency, we first quantified within-day consistency, the average correlation between each lap’s spatial activity and all other laps within the same day (**Fig. 3A**).

**Figure 3:**
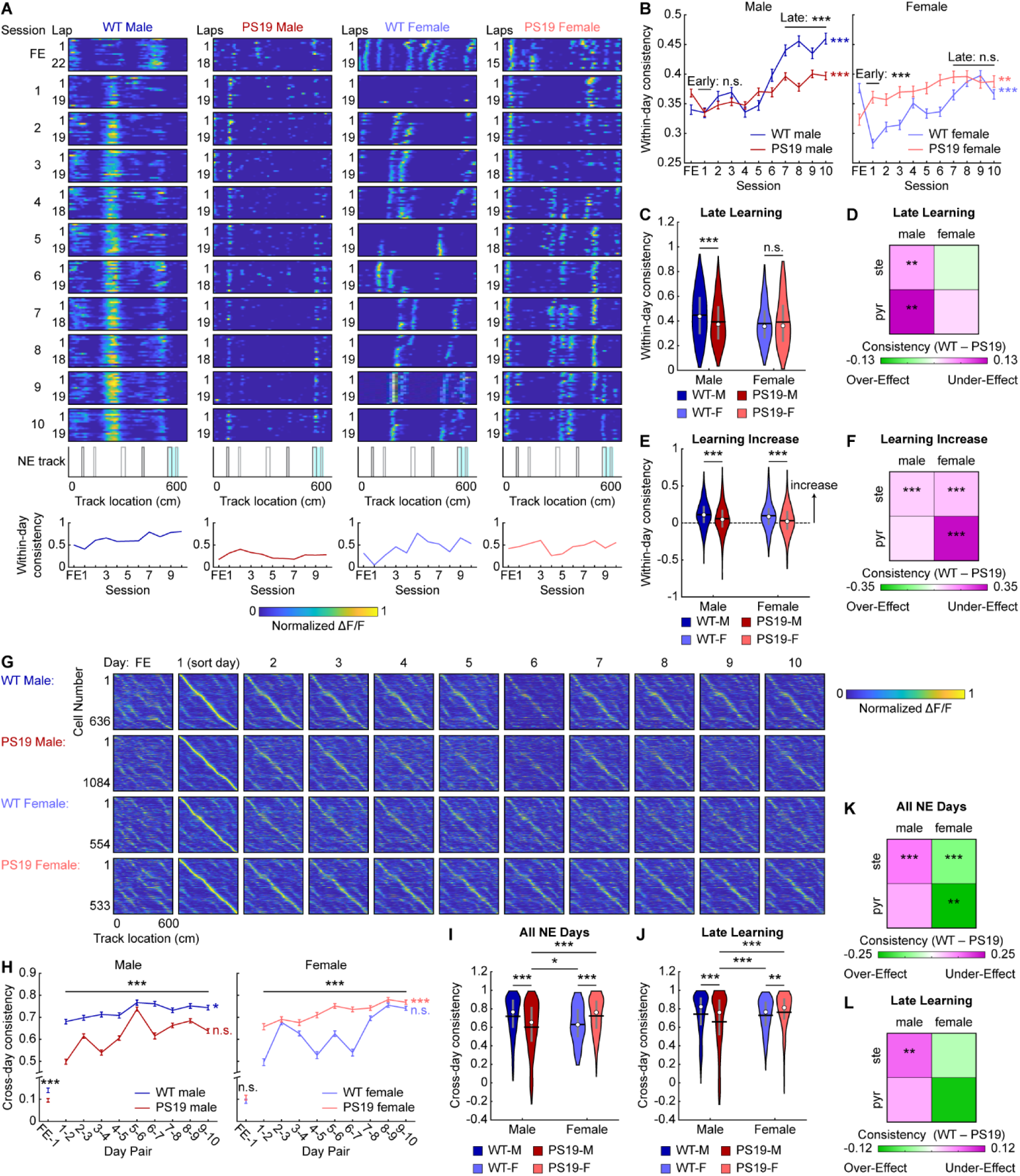
Male PS19 mice have impaired MEC within-day and cross-day activity consistency. **A.** Top: Calcium activity matrices of example cells as a function of lap and track location for the FE day and all 10 NE days for each sex/genotype category (from left to right: WT male, PS19 male, WT female, PS19 female). Middle: Environment template. Black and grey rectangles represent left and right cues, respectively. Blue rectangle represents reward zone. Bottom: Within-day activity consistency for the example cells. **B.** Within-day activity consistency across learning in male (left) and female (right) mice. Horizontal gray bars indicate p values for the group difference (early learning: unpaired Student’s t-test, late learning: linear mixed-effects model). P-values to the right indicate significant Pearson correlation of the mean value with respect to time. **C.** Within-day activity consistency averaged across late learning (days 7-10). Unpaired Student’s t-test. WT-M: WT Male, PS19-M: PS19 Male, WT-F: WT Female, PS19-F: PS19 Female **D.** Heatmap summarizing the within-day activity consistency difference between WT and PS19 mice by sex and cell type. Color represents the mean difference between WT and PS19 values averaged across late learning as in **Fig. S5C**. Unpaired Student’s t-test. **E.** The increase in within-day activity consistency between early learning (day 1) and late learning. Unpaired Student’s t-test. **F.** Heatmap summarizing the within-day activity consistency difference between WT and PS19 mice by sex and cell type. Color represents the mean difference between WT and PS19 values for the increase between late and early learning as in **Fig. S5F**. Unpaired Student’s t-test. **G.** Activity matrices for the FE day and all 10 NE days for each sex/genotype category (from top to bottom: WT male, PS19 male, WT female, PS19 female). The normalized mean activity of individual cells (each row of an activity matrix) were sorted based on the peak locations on day 1. **H.** Cross-day activity consistency across learning in male (left) and female (right) mice. Horizontal gray bars indicate p values for the group difference (linear mixed-effects model). FE-1 comparison uses unpaired Student’s t-test. P-values to the right indicate significant Pearson correlation of the mean values with respect to time. **I, J.** Cross-day activity consistency averaged across all learning days (**I**) and late learning (**J**). Unpaired Student’s t-test with Bonferroni-Holm correction. **K, L.** Heatmaps summarizing the cross-day activity consistency difference between WT and PS19 mice by sex and cell type. Color represents the mean difference between WT and PS19 values averaged across all learning days (**K**) or late learning days (**L**) as in **Fig. S5I** and **Fig. S5J**, respectively. Unpaired Student’s t-test. \**p* ≤ 0.05, \*\**p* ≤ 0.01, \*\*\**p* ≤ 0.001. In violin plots, horizontal line represents mean, white circle represents median, and whiskers represent interquartile range. Error bars in line plots represent mean ± sem. Statistical details can be found in **Table S1**. See also **Fig. S5**.

During early learning, male PS19 mice showed within-day consistency comparable to WT mice (**Fig. 3B)**, but by the late learning phase, their consistency was significantly lower (**Figs. 3B, C**), indicating a failure to stabilize their activity spatial maps over time. While lower consistency was present in both stellate and pyramidal cells, the late-learning difference was most pronounced in pyramidal cells (**Figs. 3D, S5A-C**), indicating a cell-type-specific deficit. Notably, this consistency difference was similar across grid and non-grid cells (**Figs. S5D, E**). In contrast, PS19 female mice exhibited higher within-day consistency than WT females during early learning overall and across both stellate and pyramidal cells, with differences diminished by late learning (**Figs. 3B-D, S5A-C**). Interestingly, this effect was stronger in grid than non-grid cells (**Figs. S5D, E**). Thus, impaired late-learning within-day map stability was a specific deficit to male PS19 mice.

Previous work showed that learning-related increases in within-day consistency accompany robust behavioral improvement, highlighting the importance of flexible updating spatial maps during successful learning^49^. However, both PS19 male and female mice exhibited smaller increases in within-day consistency across learning than WT mice (**Figs. 3B, E**), a pattern observed in both stellate and pyramidal cells (**Figs. 3F, S5A, B, F**). This restricted dynamic range indicates reduced flexibility of within-day spatial representations in PS19 mice despite repeated environmental exposure.

We next examined cross-day activity consistency, defined as the correlation of each cell’s lap-averaged spatial activity between adjacent days (**Fig. 3G**). Consistency between FE and NE day 1 was near-zero for all groups, with PS19 males exhibiting even lower values, indicating preserved remapping ability in PS19 mice (**Fig. 3H**). Across learning, male PS19 mice showed lower consistency than WT males (**Figs. 3H, I**). In contrast, similar to the within-day consistency, PS19 females had higher early-learning cross-day consistency than WT females, which converged by late learning (**Fig. 3H**). By days 7–10, WT males, WT females, and PS19 females reached high cross-day consistency, whereas PS19 males remained significantly lower (**Figs. 3H, J**). These differences were consistent across molecular and functional cell types (**Figs. 3K, L, S5G-L**), indicating a global deficit in cross-day consistency of PS19 males.

Together, these results show that PS19 mice exhibit impaired spatial activity consistency during learning. Male PS19 mice failed to maintain stable spatial maps both within and across days, with the within-day deficit most pronounced in pyramidal cells. Moreover, both sexes displayed inflexibility in updating spatial representations with increased experience, which may lead to inefficient learning. The more severe deficits in male PS19 mice likely contribute to their pronounced spatial learning impairments.

### Male PS19 mice do not develop a cognitive map shared by other mouse groups

The above results revealed that male PS19 mice exhibited more severely impaired cross-day activity consistency than WT males, WT females, or PS19 females. To further investigate this deficit, we considered two possibilities. First, PS19 males may have failed to develop a common “correct” cognitive map shared by other groups. Alternatively, rather than converging on a common map, each animal might develop an individual cognitive map whose internal consistency, rather than its specific pattern, supports successful spatial learning.

To distinguish these possibilities, we correlated spatial maps between individual mice (inter-mouse consistency) (**Fig. 4A**). For each mouse, a late-learning map representing the outcome of NE learning was generated by averaging the spatial activity of all common cells across days 7 to 10 (**Figs. 4A, B, S6A**). Correlations were then computed within the same sex/genotype group (within-group comparisons) or between different groups (between-group comparisons) (**Figs. 4A, C**). Within-group comparisons showed that PS19 males—whether using all common cells, pyramidal cells, or stellate cells—exhibited variability similar to WT groups, whereas PS19 females had more consistent maps (**Figs. 4D, E, S6B-E**). Thus, within-group variation does not account for the learning differences observed between PS19 males and other groups.

**Figure 4:**
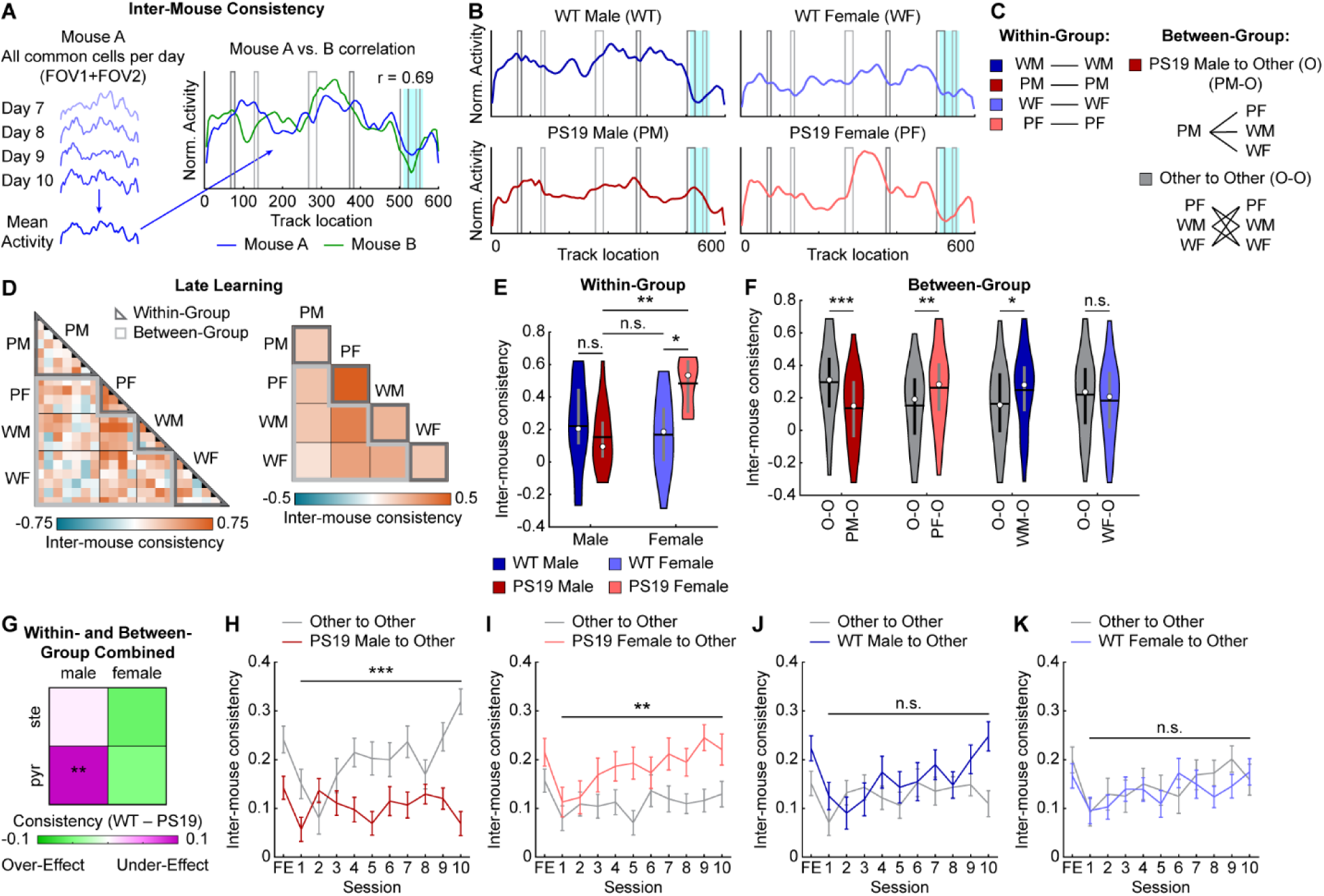
Male PS19 mice do not develop a cognitive map shared by other mouse groups. **A.** Schematic of late-learning (NE days 7-10) inter-mouse consistency calculation. **B.** Group averages of late-learning spatial activity maps as calculated in **A**. **A, B.** Black and grey rectangles represent left and right cues, respectively. Blue rectangle represents reward zone. **C.** Schematic for within-group (left) and between-group (right) comparisons. **D.** Left: Pairwise inter-mouse consistency sorted by genotype/sex group. Right: Average inter-mouse consistency for each group-to-group block. **E.** Pooled inter-mouse consistency for pairwise within-group comparisons. Unpaired Student’s t-test with Bonferroni-Holm correction. **F.** Pooled inter-mouse consistency for pairwise between-group comparisons with each genotype/sex group serving as the reference group as indicated. The Unpaired Student’s t-test. **G.** Heatmap summarizing the inter-mouse consistency difference between WT and PS19 mice by sex and cell type. Color represents the mean difference between WT and PS19 values from **Fig. S6B, C**. All pairwise correlations between mice in a given group with all mice were pooled for calculating the difference. **H-K.** Inter-mouse consistency for between-group comparisons across learning using PS19 males (**H**), PS19 females (**I**), WT males (**J**), and WT females (**K**) as a reference group. Horizontal gray bars indicate p values for the group difference (linear mixed-effects model). \**p* ≤ 0.05, \*\**p* ≤ 0.01, \*\*\**p* ≤ 0.001. In violin plots, horizontal line represents mean, white circle represents median, and whiskers represent interquartile range. Error bars in line plots represent mean ± sem. Statistical details can be found in **Table S1**. See also **Fig. S6**.

In contrast, PS19 male maps were less similar to those of other groups. Between-group comparisons using common cell maps revealed significantly lower PS19-male-to-Other consistency compared with Other-to-Other consistency, indicating that PS19 male maps differed from all other groups (**Figs. 4D, F)**. This effect was specific to pyramidal but not stellate cells **(Figs. S6B, C, F, G**) and was absent in other sex/genotype groups (**Figs. 4D, F, S6B, C, F, G).** When combining all within- and between-group comparisons, only pyramidal cells in PS19 males showed significantly lower inter-mouse consistency relative to WT mice (**Fig. 4G**). These findings suggest that while other mice converged on a common cognitive map by the end of learning, PS19 males developed distinct maps, with deficits primarily affecting pyramidal cells.

We next asked whether the cognitive map differences in PS19 males were present at the onset of learning or emerged gradually. To address this, we tracked map development across days by calculating inter-mouse consistency between groups on each learning day. While PS19-male-to-Other consistency remained low, Other-to-Other consistency steadily increased over learning and diverged progressively from PS19-male-to-Other consistency (**Fig. 4H**). This pattern was not observed when substituting any other sex/genotype group for PS19 males (**Figs. 4I-K**). These findings suggest that although all groups began with similarly variable maps, PS19 males uniquely failed to develop the shared cognitive map that emerged across other groups during learning.

In summary, successful spatial learning requires both high internal consistency within each animal’s cognitive map and convergence toward a common, “correct” map. The failure of PS19 males to form this shared map, primarily in pyramidal cells, likely contributes to their spatial learning deficits, underscoring the critical role of proper cognitive map formation in spatial memory.

### Spatial activity of male PS19 mice is anchored to visual cues

Following the above finding, we asked what kind of map PS19 males formed during learning and how it differed from other groups. The impaired speed modulation observed in PS19 males (**Figs. 2J-L**) and low spatial activity consistency (**Figs. 3B-D**) suggest disrupted path integration, whereby self-motion cues (e.g., running speed and acceleration) are integrated over time to estimate position independently of external landmarks^40,94–96^. Disruption of this process would bias spatial representations toward cue-rich regions and weaken representations in cue-lacking regions^54,97^ (cue anchoring; **Fig. 5A**). Consistent with this prediction, we observed a visual trend toward cue anchoring in PS19 males in both group-averaged maps (**Fig. 4B**) and individual late-learning maps (**Fig. S6A**).

**Figure 5:**
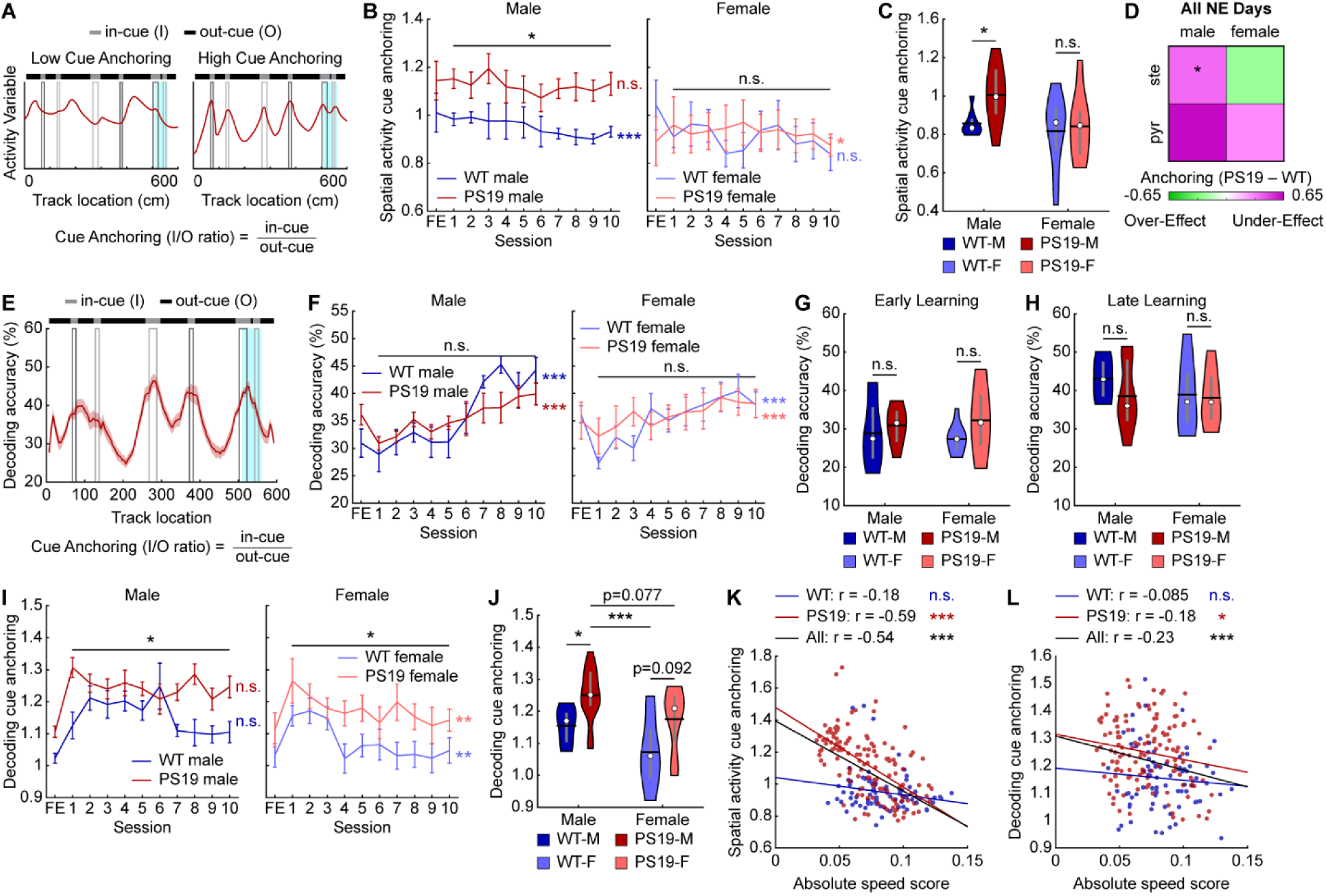
Spatial activity of male PS19 mice is anchored to visual cues. **A.** Examples of spatial activity with low (left) and high (right) cue anchoring and the cue anchoring calculation. Vertical black and grey rectangles represent left and right cues, respectively. Blue rectangle represents reward zone. **B.** Spatial activity cue anchoring across learning in male (left) and female (right) mice. **C.** Spatial activity cue anchoring averaged across all learning days. WT-M: WT Male, PS19-M: PS19 Male, WT-F: WT Female, PS19-F: PS19 Female **D.** Heatmap summarizing the spatial activity cue anchoring difference between PS19 and WT mice by sex and cell type. Color represents the mean difference between PS19 and WT values averaged across all learning days as in **Fig. S7F**. Unpaired Student’s t-test. **E.** Example of decoding accuracy as a function of track position. From PS19 males averaged across the 10 learning days. Vertical black and grey rectangles represent left and right cues, respectively. Blue rectangle represents reward zone. **F.** Full track decoding accuracy across learning in male (left) and female (right) mice. **G, H.** Full track decoding accuracy averaged across early (day 1, **G**) and late learning (days 7-10, **H**). **I.** Decoding cue anchoring across learning in male (left) and female (right) mice. **J.** Decoding cue anchoring averaged across all learning days. Unpaired Student’s t-test with Bonferroni-Holm correction. **K, L.** Correlation between absolute speed score and spatial activity cue anchoring (**K**) or decoding cue anchoring (**L**) in male mice. Each point represents the activity from one FOV from one session. Pearson correlation. \**p* ≤ 0.05, \*\**p* ≤ 0.01, \*\*\**p* ≤ 0.001. In violin plots, horizontal line represents mean, white circle represents median, and whiskers represent interquartile range. Violin plots use unpaired Student’s t-test. Error bars and shaded regions in line plots represent mean ± sem. In line plots, horizontal gray bars indicate p values for the group difference (linear mixed-effects model), and p-values to the right indicate significant Pearson correlation of the mean value with respect to time. Statistical details can be found in **Table S1**. See also **Figs. S7, S8**.

To quantify cue anchoring, we calculated the ratio of activity in in-cue (I) versus out-cue (O) regions (spatial activity cue anchoring) (**Fig. 5A**). PS19 males showed a trend of higher in-cue activity but similar out-cue activity compared to WT males (**Figs. S7A-C**), resulting in significantly higher cue anchoring across learning (**Figs. 5B, C**). In contrast, PS19 females exhibited insignificant trends of elevated activity in both in-cue and out-cue regions (**Figs. S7A-C**), yielding comparable cue anchoring to WT females (**Figs. 5B, C**). Notably, cue anchoring in WT males decreased across learning (**Fig. 5B**), suggesting a shift toward a more global spatial representation that might support successful learning. However, cue anchoring of PS19 males failed to improve, suggesting a deficit. When split by molecular cell type, pyramidal cells of male PS19 mice showed the highest cue anchoring (**Figs. S7D-F**) and the largest difference from WT males (**Fig. 5D**). For functional cell types, both grid and non-grid cells in male PS19 mice showed a trend of stronger cue anchoring than WT males (**Figs. S7G-I**).

We further validated activity cue anchoring using population decoding of track positions^49,98^. A decoder built from population activity in odd laps at individual 5-cm spatial bins was used to predict positions from even-lap activity, with accuracy defined as the percentage of correctly decoded positions within ±2 bins from the original bin. Because decoding accuracy is cell-number dependent^49,98^, decoding was performed with 100 random samples of 30 cells for each field-of-view (FOV). All cells were included because many FOVs lacked sufficient numbers of common, stellate, or pyramidal cells for separate analyses.

Decoding accuracy was assessed across the full track and by location (**Fig. 5E**). Consistent with our previous work^68^, WT mice exhibited gradually improved accuracy during learning (**Fig. 5F**). PS19 males exhibited lower late-learning accuracy than WT males, while female WT and PS19 mice reached similar levels (**Figs. 5F-H**). Both PS19 males and females exhibited trends of higher accuracy at in-cue regions than WT mice during early learning and lower accuracy at out-cue regions during late learning (**Figs. S7J-L**), resulting in higher ratios of decoding accuracy at in-cue versus out-cue regions (decoding cue anchoring) throughout learning (**Figs. 5E, I, J**). Notably, PS19 males showed the highest anchoring among all groups (**Fig. 5J**). Cue anchoring was more pronounced in grid than non-grid cells (**Figs. S7M-O**). Across learning, both in-cue and out-cue decoding improved (**Figs. S7J, K**), but cue anchoring declined (**Fig. 5I**), supporting the idea that reduced cue anchoring favors successful spatial learning and indicating that elevated anchoring in PS19 mice reflects an activity deficit.

Together, these results demonstrate that PS19 mice, especially males, exhibited pronounced cue anchoring in MEC spatial representations. Cue anchoring was stronger in grid than non-grid cells and in pyramidal than stellate cells. The relatively weak representation of non-cue regions could stem from reduced speed cell number and activity speed modulation (**Figs. 2H-L**), both of which are essential for providing self-motion signal during path integration when external landmarks are unavailable^6,99^. This connection is supported by a strong negative correlation between absolute speed score and cue anchoring in PS19, but not WT, males (**Figs. 5K, L**), indicating that poor speed modulation is robustly associated with higher cue anchoring specifically under conditions of behavioral impairment. This relationship was observed across cell types but was strongest in pyramidal and non-grid cells (**Figs. S8A-F**). Thus, impaired self-motion integration in PS19 males likely drives excessive reliance on cue regions rather than the formation of a coherent, global cognitive map. These deficits are potentially manifested as path integration deficits, which have been observed in PS19 mice and are associated with pTau accumulation in the EC^61^, ultimately leading to disrupted spatial memory formation.

### Tau accumulation and activity deficits in male PS19 mice predominantly occur in pyramidal cells

Overall, PS19 mice, particularly males, exhibited widespread deficits across multiple activity domains, including general activity features (activity level, absolute speed score, and spatial selectivity), map consistency (within-day consistency, cross-day consistency, and inter-mouse consistency), and global spatial representation (spatial activity cue anchoring and decoding cue anchoring) (**Fig. S9**). Given the stronger pTau accumulation in pyramidal cells than in stellate cells (**Figs. 1, S1,** summarized in **Fig. 6A)** and the differences in the activity deficits across these cell types, we examined whether the imbalance in pTau accumulation corresponds to the severity of activity deficits, a key step in linking tauopathy to spatial memory impairment.

**Figure 6:**
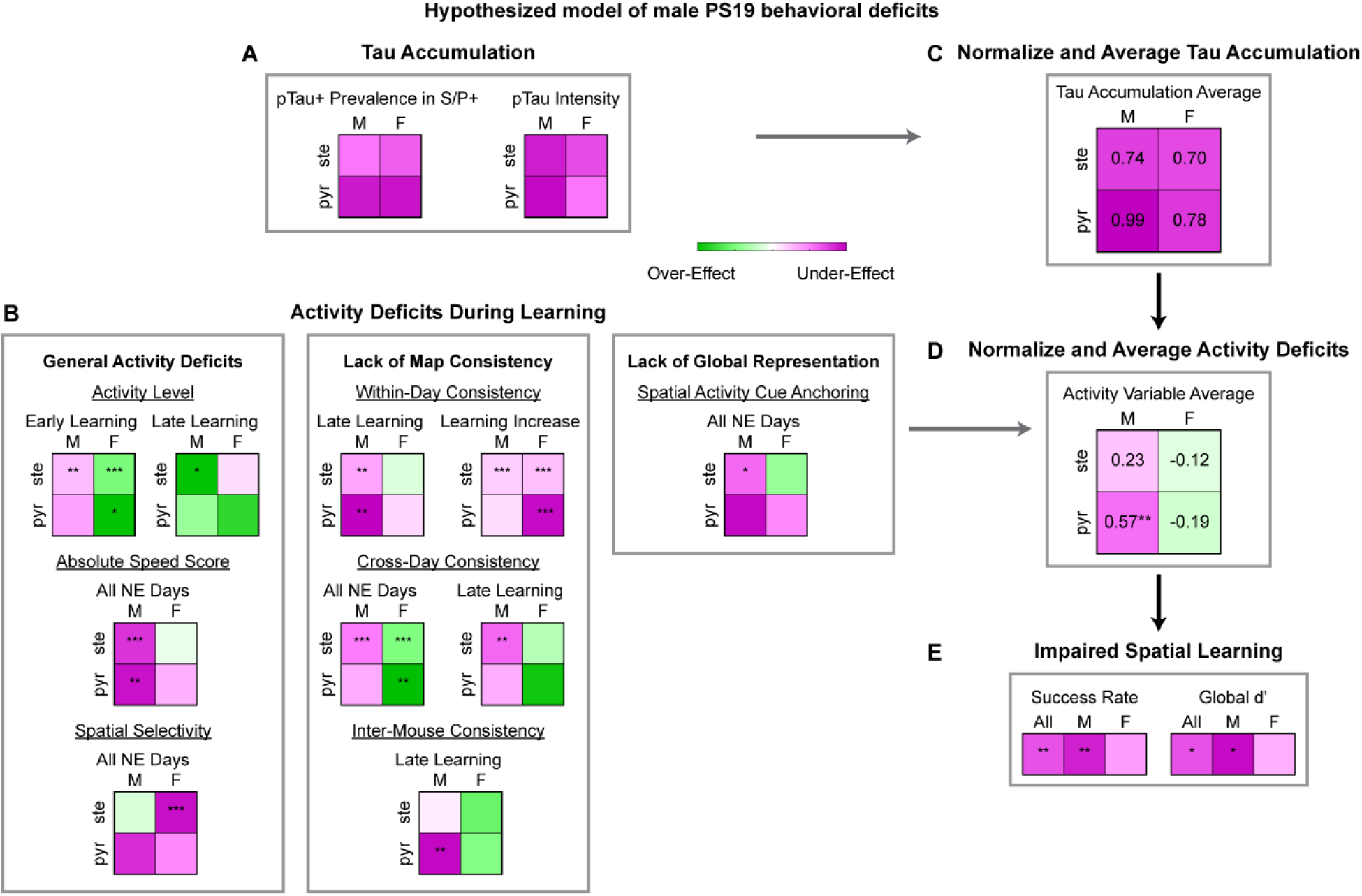
Tau accumulation and activity deficits in male PS19 mice predominantly occur in pyramidal cells. **A-E.** Summary of potential mechanism explaining PS19 behavioral deficits, whereby tau accumulation, particularly in pyramidal cells, leads to changes in MEC activity, which leads to impaired spatial learning. **A, B, E.** All heatmaps summarizing the sex (male, M vs. female, F) and cell type (stellate, ste, S vs. pyramidal, pyr, P) split of tau accumulation (**A**), activity deficits (**B**), and behavioral deficits (**E**) are identical to those presented in earlier figures. **C, D.** Normalized and averaged 2x2 heatmap of tau accumulation (**C**) and activity deficits (**D**) showing overall tau accumulation and activity deficits are most prominent in pyramidal cells of male PS19 mice. Individual activity heatmaps were normalized to their maximum absolute value, so the individual heatmap values and the color range for the average heatmaps range between -1 and 1. One-sample Student’s t-test for difference from 0. Note that the average heatmaps should only be interpreted qualitatively. \**p* ≤ 0.05, \*\**p* ≤ 0.01, \*\*\**p* ≤ 0.001. Statistical details can be found in **Table S1**. See also **Fig. S9**.

To provide a qualitative measure of deficit severity, we summarized the 2×2 heatmaps capturing cell-type– and sex-specific differences between PS19 and WT mice across all activity measures (**Figs. 2-5, S4-S7,** summarized in **Fig. 6B)**. Tau and activity heatmaps were normalized by their maximum absolute values, yielding values between −1 and 1, and averaged to generate summary heatmaps for tau accumulation and overall activity differences between WT and PS19 mice (**Figs. 6C, D**). PS19 males exhibited strong activity under-effects, particularly in pyramidal cells, reflecting pronounced activity deficits below WT levels (**Fig. 6D**), consistent with their higher pTau burden in pyramidal cells (**Fig. 6C**) and potentially leading to severe spatial learning impairments (**Fig. 1, summarized in Fig. 6E**). In contrast, PS19 females showed relatively mild over-effects, especially in pyramidal cells. This trend of exceeding of WT level may also contribute to cognitive disruption (e.g., hyperactivity^100^), albeit to a lesser degree, likely reflecting their lower pTau burden (**Fig. 6C**) and resulting in mildly impaired spatial learning (**Figs. 6E**).

Together, these findings suggest that selective tau accumulation in pyramidal cells is closely associated with activity dysfunction and spatial memory impairment, establishing a link between tauopathy and cognitive deficits.

### MEC activity changes are predictive of behavioral deficits in PS19 mice

While the above qualitative analyses suggest links between tau burden, MEC activity impairments, and spatial learning deficits, we further strengthen these relationships using quantitative modeling to identify which activity variables best predict learning performance and how cell types with different tau burdens contribute to these predictions.

We utilized a general linear model (GLM)^101^ to predict each mouse’s success rate in a given session based on behavior or activity variables within that session (**Fig. 7A**), using data from all NE days. Model predictions were evaluated by calculating the percentage of behavioral variance explained (R^2^), where 100% indicates a perfect match between predicted and true behaviors. Predictor variables included behavioral variables (mean velocity), general activity variables (activity level, raw speed score, absolute speed score, and spatial selectivity), map consistency variables (within-day consistency, cross-day consistency, and inter-mouse consistency), and global representation variables (spatial activity cue anchoring and decoding cue anchoring). Activity variables were derived from common cells, as described previously, except decoding cue anchoring, which used all cells. Sex and genotype were included as confounding variables. All predictor variables were standardized using z-score normalization and used in an initial model with the confounding variables. To identify the most informative predictors, variables that did not significantly contribute were iteratively removed while retaining confounding variables (**Fig. S10A**). This procedure substantially increased variance explained compared to shuffled predictions, where success rates were randomized to disrupt correlations with predictors^102^ (**Figs. S10A, B**).

**Figure 7:**
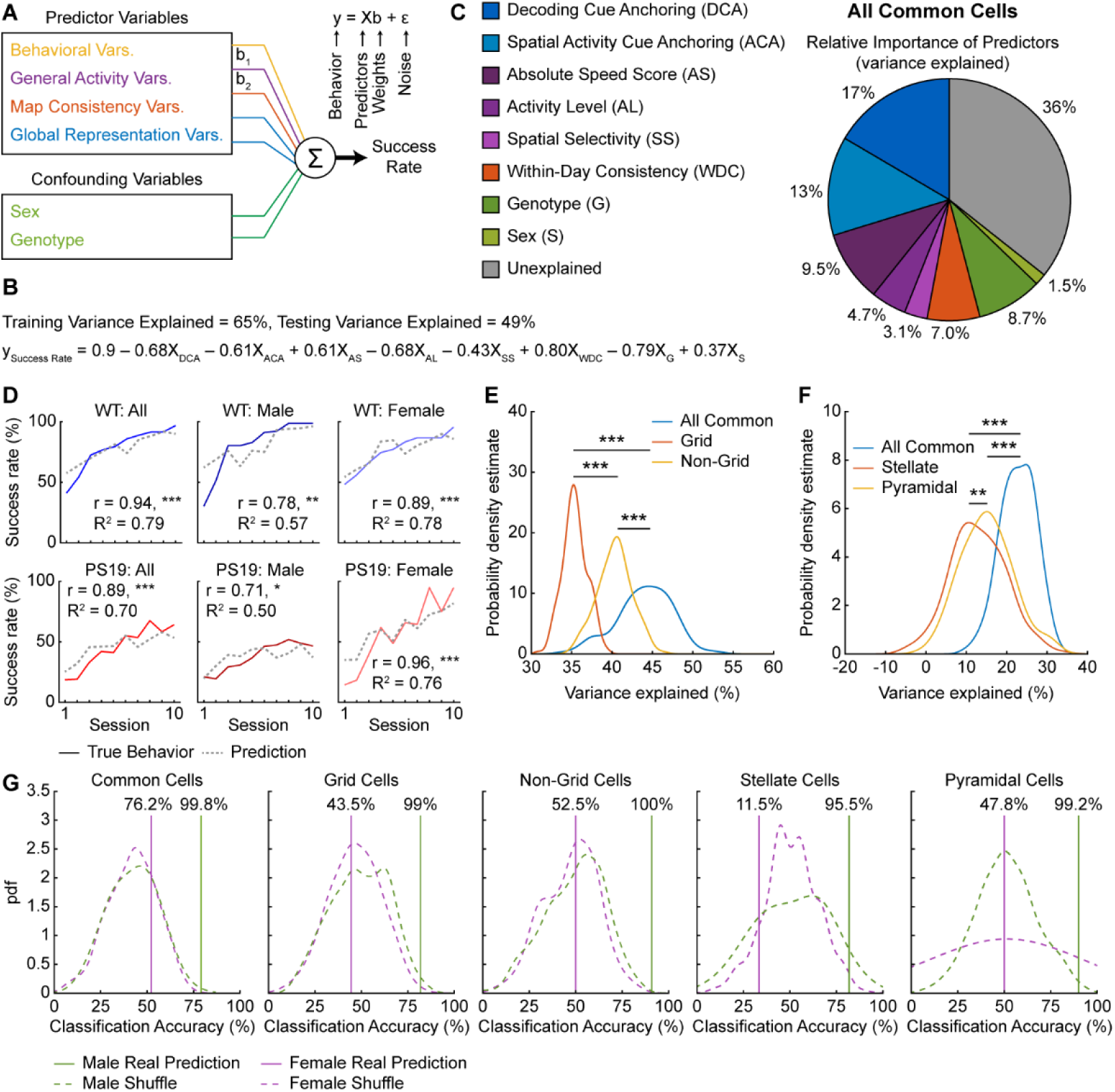
MEC activity changes are predictive of behavioral deficits in PS19 mice. **A.** Schematic showing use of a general linear model (GLM) to predict behavior using behavioral, general activity, map consistency, global representation, and confounding variables. **B.** Final LOOVC model performance. Training and testing variance explained are calculated by pooling LOOCV iterations. Model equation is calculated by averaging coefficients from each iteration. Each X represents the value of a given predictor or confounding variable. **C.** Relative contribution of predictor and confounding variables to behavioral variance explained in a GLM using activity from all common cells. Colors correspond to variable categories in **A**. **D.** Comparison of mean success rate (colored lines) and model predictions (dashed grey lines) across learning for WT (top) and PS19 (bottom) in all (left), male (middle), and female (right) mice using activity from all common cells. Pooled test predictions across all LOOCV iterations were used to calculate prediction means. Pearson correlation. **E, F.** Probability distribution of variance explained for models (**E**: all-common-cell, grid cell, non-grid cell; **F**: all-common-cell, stellate cell, pyramidal cell) based on subsampled activity data. Each mouse was randomly subsampled 200 times, such that the activity variables of all three groups were based on identical numbers of cells. Unpaired Student’s t-test. **E**: common vs. grid: d (Cohen’s D) = 3.2. common vs. non-grid: d = 1.3. grid vs. non-grid: d = 2.7. **F**: common vs. stellate: d = 1.7. common vs. pyramidal: d = 1.4. stellate vs. pyramidal: d = 0.31. **G.** Classification accuracy of genotype based on modeling of mouse behavior from all common, grid, non-grid, stellate, and pyramidal cells (from left to right). Vertical line labels indicate the percentile of the real prediction classification among the shuffles. \**p* ≤ 0.05, \*\**p* ≤ 0.01, \*\*\**p* ≤ 0.001. Statistical details can be found in **Table S1**. See also **Figs. S10 and S11**

The final model retained the six activity variables that were never removed across all iterations: activity level, absolute speed score, spatial selectivity, within-day consistency, spatial activity cue anchoring, and decoding cue anchoring (**Figs. 7B, S11A**). Individually removing any of these variables reduced variance explained, whereas adding back excluded variables had little effect, highlighting the importance of the final variables (**Fig. S10C**). Excluding sex and genotype during variable selection did not alter the final activity variables, indicating no bias from confounding variables (**Fig. S10D**). Correlation analyses confirmed that these activity variables were strongly associated with success rate (**Figs. S10E-N**), with four (absolute speed score, within-day consistency, decoding cue anchoring, spatial activity cue anchoring) significantly correlated within each genotype (**Figs. S10F, J, M, N**).

Dominance analysis quantified each variable’s contribution by measuring the effect of removing it^103^ (**Fig. 7C**). The six activity variables collectively explained 54% of the behavioral variance, while confounding variables accounted for only 10% (**Fig. 7C**), underscoring the strong link between MEC activity and learning. Predicted success rates closely matched the true learning curves across groups, showing significantly high correlations (r) and variance explained (R^2^) (**Fig. 7D**), though individual mice showed some variability (**Fig. S11B**).

Given that grid cells play a central role in MEC spatial coding^104–107^ and are disrupted in tauopathy, as reported previously^9,39,40^ and here, we tested whether their activity is particularly predictive of behavior by comparing grid, non-grid, and all common cells (including grid and non-grid cells together). Because using a greater number of cells may increase predictive ability^108^, we subsampled equal numbers of cells per category and repeated the prediction 200 times using the final model variables to generate distributions of variance explained. While subsampling slightly reduced variance explained, the all-common-cell model outperformed both grid and non-grid models, and non-grid cells explained more variance than grid cells (**Fig. 7E**). Effect sizes (Cohens d^109^) were large and only a few subsamples were needed to reach 80% power^109^ (80% probability of obtaining a significant result if a true difference exists) (**Figs. 7E, S11C**). Thus, both grid and non-grid cell activity contribute to predicting behavior, with non-grid activity being more informative.

Because male PS19 mice show greater activity impairment in pyramidal than stellate cells, we tested whether pyramidal activity better predicts spatial learning. Using all common cells and either stellate or pyramidal cells from the common cell population, we applied the same subsampling procedure to select equal numbers per category. The all-common-cell model explained more variance than either cell-type model, and pyramidal cells outperformed stellate cells (**Fig. 7F**). Effect sizes were large for common versus cell-type models and moderate for stellate versus pyramidal cells (**Fig. 7F**). Only few subsamples were needed to achieve 80% power for common versus cell-type comparisons, but more were required for stellate versus pyramidal cells (**Fig. S11D**). These results show that both cell types contribute to predicting learning, with pyramidal cells being more informative.

Finally, we tested whether these activity variables could classify mouse genotype. Using the final model variables—excluding sex and genotype as confounding variables and without subsampling—we averaged late-learning predictions for each mouse, which was further classified as WT or PS19 based on proximity of the prediction to group averages of real behaviors. Male classification exceeded chance across all cell types (all common cells, grid, non-grid, stellate, and pyramidal cells) (**Fig. 7G**), indicating that these variables captured PS19 behavioral deficits regardless of functional or molecular cell type. Female classification was at chance (**Fig. 7G**), likely reflecting minimal deficits of PS19 females.

In summary, MEC activity—including in grid, non-grid, stellate, and pyramidal cells—predicts spatial learning. Non-grid and pyramidal cells contributed to better prediction than grid and stellate cells, respectively. Altered activity patterns in PS19 mice—encompassing activity level, spatial selectivity, speed modulation, map consistency, and cue anchoring—likely underlie their spatial learning deficits, which is further supported by their ability to classify genotype.

## DISCUSSION

The link between EC neural activity and spatial memory deficits in tau pathology in aging and early AD remains poorly understood. To investigate this, we performed two-photon calcium imaging of layer 2 excitatory neurons in the MEC of WT and PS19 mice as they learned to navigate a novel VR environment for 10 days. pTau accumulation in MEC layer 2 increased markedly after 6 months, particularly in pyramidal cells. Interestingly, despite largely comparable pTau accumulation across sexes, only PS19 males exhibited severe spatial learning deficits, whereas females were less affected. To identify underlying neural mechanisms, we quantified a broad set of behavioral and activity features and used modeling to predict learning performance and distinguish genotypes. Spatial learning deficits in PS19 mice were primarily associated with impaired speed modulation, reduced spatial map stability, and overrepresentation of cue-rich versus cue-poor regions. These observations suggest deficits in path integration, which could prevent the formation of a reliable and global cognitive map and ultimately lead to impaired spatial memory. The activity deficits were most robust in pyramidal cells, consistent with their greater pTau burden relative to stellate cells. Furthermore, learning performance was well predicted by MEC activity, with higher accuracy by non-grid than grid cells and by pyramidal than stellate cells. MEC activity also reliably distinguished male PS19 mice from WT controls. Together, these results strongly link tau pathology to MEC dysfunction and highlight specific neural activity features that likely drive spatial learning deficits. These features may inform future diagnostic biomarkers for early AD and guide therapeutic strategies for aging- and AD-related tau pathology by focusing research on EC circuit function.

### Choice of mouse models

Our study used the PS19 mouse model, which overexpresses the human 1N4R tau P301S mutant under the mouse prion promoter^58^. While this overexpression model does not fully recapitulate tau pathology in aging and early AD, it offers several advantages. First, high tau expression in MEC neurons induces robust activity changes, allowing us to examine the relationship between tau pathology and neuronal function. Second, PS19 mice develop spatial memory deficits early, before substantial neurodegeneration occurs. This temporal window allows more meaningful comparison with tau pathology in aging and early AD, and facilitates single-neuron analyses without major confounds from widespread network disruption. Third, memory impairments arise prior to paralysis, allowing mice to perform VR-based navigation tasks that require sustained locomotion. Finally, the absence of amyloid-β in this model may recapitulate aspects of human aging, as tau accumulation is almost universal at old age, even in the absence of amyloid-β^26^. Thus our study potentially offers insights into human aging not offered by standard mouse aging studies, as mice do not naturally develop tau accumulations in old age^110^. Although tau expression in P19 mice is not restricted to the MEC, we find that MEC pTau accumulation patterns strongly correlate with MEC neural activity features, which in turn predict spatial learning and reliably distinguishes PS19 from WT mice. Together, these findings establish a link between tau pathology, MEC dysfunction, and spatial memory deficits.

In contrast, other tauopathy models are less suitable for these goals. EC-tau mice^111–113^, expressing human P301L tau mutant selectively in entorhinal layer 2/3, show spatial memory deficits only at ∼30 months^39,111^, with substantial neuronal loss beginning at 24 months^39,113^. These features limit VR navigation due to markedly reduced locomotion in aged mice^114^ and confound neural activity interpretation. In addition, the severe neuronal loss in this model compares poorly with the pathology observed in aging and early AD. Human MAPT knock-in (KI) mice, expressing human WT tau isoforms or mutant human tau^115–119^, avoid overexpression artifacts but often fail to develop robust tau pathology or spatial cognitive deficits even at advanced ages^116–118,120,121^. Robust phenotypes typically require combining multiple MAPT mutations^122,123^, crossing with Aβ/APP mouse models^119,124^, or seeding with human brain derived tau^125^. These manipulations complicate experiments using crosses with Thy1-GCaMP6 mice^67^, which ensure stable GCaMP6f expression for long-term monitoring of neural activity during learning but could alter the development or timing of tau-pathology-related phenotypes. Moreover, learning impairments in these models often coincide with severe synaptic loss and neurodegeneration^122,123^, again making them difficult to compare with aging and early AD and hindering efforts to isolate tau-induced changes in neural activity from the effects of synaptic or neuronal loss.

### Comparison with previous studies on the MEC and the hippocampus

Although many studies have reported a wide range of malfunctions of MEC neurons in AD and tauopathy models^39–42^, most have focused on neural activity at a single time point of navigation. Because spatial learning and memory depend on repeated exploration across days^47,48^, it has remained unclear which specific MEC activity features drive learning deficits. By longitudinally tracking MEC neural activity alongside behavior over 10 days of learning, we identify key activity features that mechanistically link tau pathology to impaired spatial memory formation.

#### Impaired path integration potentially leads to unsuccessful spatial memory in PS19 mice

Among aberrant activity features reported previously^39–42^ and in our study, a prominent predictor of learning performance was the ability of the MEC map to gradually stabilize into a coherent global representation. PS19 males showed reduced stabilization of spatial representations, consistent with our prior finding that MEC spatial consistency is necessary for successful learning^49^. This instability parallels observations in 3xTg mice, which model both amyloid-β and tau pathology^126^, in which spatial consistency of hippocampal place cells likewise fails to increase across learning^127^.

In addition to instability, MEC spatial representations of PS19 mice were biased toward cue-rich regions, with weaker encoding of non-cue areas. This bias was accompanied by reduced speed modulation and a lower proportion of speed-modulated MEC neurons, features that are critical for path integration based on self-motion signals independent of external landmarks^6,99^. Consistent with this interpretation, PS19 mice showed a strong association between impaired speed encoding and excessive cue anchoring, suggesting that disrupted path integration prevents the formation of a coherent, global spatial map. Supporting this view, impaired path integration has previously been reported in PS19 mice and was associated with pTau accumulation in the EC^61^. The degree of cue anchoring and impairment in speed modulation was more pronounced in pyramidal cells, which exhibited the strongest pTau accumulation, suggesting a cell-autonomous effect of pTau on these activity deficits. Additionally, such deficits may partly arise from dysfunction of the locus coeruleus (LC)^61,128–130^, which also exhibits robust tau burden in PS19 mice^128,129^ and sends noradrenergic projections to multiple brain regions, including the MEC, to control motor function, attention, and memory^131,132^. Consistent with our observations, reduced speed modulation has also been reported in rTg4510 tauopathy mice^40^.

Although the cellular mechanisms cannot be directly assessed in humans, impaired navigation based on self-motion cues and deficits in path integration have been consistently observed in older adults^133,134^ and in individuals in early AD^135–137^. Notably, the LC is the first site showing abnormally phosphorylated pretangle tau, preceding the onset of prion-like seeding activity of abnormal tau in the EC^3,26,138,139^, and LC neurons show progressive degeneration that correlates with cognitive decline^140^. These findings are consistent with a contributory role of LC dysfunction in the observed path integration deficits.

Together, these convergent findings support a model in which impaired self-motion computation in AD promotes overreliance on external landmarks, disrupts path integration, destabilizes spatial representations, and ultimately impairs spatial memory formation.

#### Inflexibility of MEC maps in PS19 mice

We observed reduced experience-dependent flexibility of MEC map dynamics in PS19 mice, as both male and female mice showed minimal changes in map consistency within days, whereas WT mice displayed the expected initial disruption followed by progressive stabilization. WT mice also exhibited increased speed modulation as learning proceed, but PS19 males failed to show the change and remained at a low level, reflecting an inflexibility of neural activity with respect to speed over the course of learning. Similar hippocampal inflexibility has been reported in APP mice, where neuronal firing patterns remained unchanged throughout learning, whereas WT cell activity became increasingly task-specific^141^. This neural rigidity likely impairs adaptation to novel environments and may reflect tau-pathology-related deficits in synaptic plasticity^142^.

#### Widespread entorhinal circuit disruption in PS19 mice

While previous studies focused primarily on grid cell dysfunction^39–42^ and reported inconsistent impairment of non-grid cells^39,40^, we found that activity deficits in PS19 grid cells extended to non-grid cells. Both populations contributed to predicting learning behavior, with non-grid cells performing even better than grid cells, and predictions based on the full ensemble surpassing either population alone. These findings support the view that accurate spatial representation arises from integrated activity across heterogeneous MEC neurons, rather than from a specialized functional subgroup^143^. Importantly, our results indicate that spatial memory impairments in tauopathy reflect widespread entorhinal circuit disruption.

#### Contributions of other brain regions

While MEC activity features explained over half of behavioral variance, residual variance likely stems from dysfunction in other regions, such as the LC discussed above. Another probable contributor is the hippocampus, which is also heavily involved in spatial cognition^6^ and accumulates phosphorylated tau and neurofibrillary tangles in PS19 mice^58^. Indeed, the hippocampal circuit in PS19 mice shows synaptic pathology, neural degeneration, altered glia dysfunction, and transcriptomic changes^58,144,145^. Moreover, tau pathology may propagate from MEC to hippocampal subfields, including the dentate gyrus and CA fields, further disrupting hippocampal circuits during spatial learning^112,113^.

### Difference in molecular cell types

#### Tau pathology is associated with impaired spatial encoding at the cellular level

We examined molecularly defined excitatory cell types in MEC layer 2—stellate and pyramidal cells—in PS19 mice. Tau pathology preferentially accumulated in pyramidal cells, which exhibited more severe activity impairments than stellate cells, including reduced spatial selectivity and speed modulation, lower activity stability, and stronger cue anchoring during learning. Given that pyramidal cells are highly speed-modulated and contribute to self-motion processing^146,147^, their selective vulnerability provides a mechanistic link between tau accumulation and impaired path integration, which may destabilize spatial representations and ultimately lead to spatial memory deficits. Accordingly, pyramidal cells in male PS19 mice exhibited higher AT8 immunoreactivity than females, paralleling their more severe learning deficits, and pyramidal cell activity better predicted learning performance across genotypes. Although *in vivo* labeling of intracellular tau in 7-10-month-old mice remains technically challenging, owing to the limited availability of well-characterized tau-specific fluorescent reporters and high brain autofluorescence in aging animals^148^, our results nonetheless provide compelling evidence for the association between tau accumulation and impaired spatial encoding at the cellular level.

Consistent with our observations, in humans, calbindin+ pyramidal cell clusters are most prominent in the caudal EC layer 2^149^, coinciding with robust pTau accumulation in the same region in very early Braak stages of AD^3,150^. At this stage, pTau and pyramidal cells also exhibit strong colocalization at the cellular level^83^. Tau in pyramidal cells of EC layer 2 can be further propagated to CA1, disrupting neural excitability and contextual learning^83^. While these findings together suggest that pyramidal cell dysfunction in EC layer 2 may underlie early EC–hippocampal circuit deficits and spatial memory impairments in early AD, other studies reported that layer 2 reelin+ stellate cells are preferentially susceptible to tau pathology^151^. Further research is needed to clarify the relative contributions of pyramidal and stellate cell dysfunction to early tau-related circuit impairments.

#### Tau pathology affects neural excitability in a cell-type-, sex-, and learning-dependent manner

Interestingly, although PS19 mice showed an increased fraction of active neurons—driven primarily by stellate cells—the activity of individual stellate and pyramidal neurons varied in a sex- and learning-dependent manner, indicating a complex and dynamic impact of tau pathology on MEC excitability. These findings extend prior reports of MEC hypoactivity^40^, including deficits in grid cells^39^ and stellate cells^41^ by revealing previously unrecognized modulation by sex and learning stage. Notably, intracellular pTau accumulation—more prominent in pyramidal cells—did not directly correlate with overall activity levels, suggesting that neuronal excitability is not solely driven by pTau or neurofibrillary tangles. Instead, altered excitability may be mediated by soluble tau species, consistent with *in vivo* two-photon imaging studies showing tau-induced hypoactivity independent of tangle formation^152,153^.

In contrast, stellate cell hyperexcitability has also been documented in APP-transgenic rats^51^, consistent with the well-established role of amyloid-β in promoting neural hyperactivity across brain regions^154^. Amyloid-β-induced neuronal hyperexcitability has been linked to plaques^155,156^, intraneuronal amyloid-β^157^, and soluble amyloid-β species^155,158^. Notably, in 3xTg-AD mice, stellate cells were hyperexcitable across ages, whereas pyramidal cell hyperactivity was restricted to younger mice^42^. These findings raise the possibility that the combined effects of amyloid-β and tau may differentially affect MEC network activity over time and across cell types compared to each of them alone.

Overall, our results suggest that both pyramidal and stellate cells contribute to tauopathy-related dysfunction, but likely via distinct mechanisms tied to their differential susceptibility to tau pathology and their unique roles in MEC circuit dynamics.

### Sex-specific behavioral and neural activity impairments

We observed pronounced sex-specific differences in MEC tau pathology and associated circuit dysfunction. Although males showed a trend toward a smaller overall pTau-positive area and similar proportions of pTau-positive stellate and pyramidal neurons compared with females, the local intensity of pTau within positive regions—particularly in pyramidal neurons—was higher in males, indicating a greater local tau burden. Correspondingly, male PS19 mice exhibited more severe spatial learning impairments and disrupted neural activity than females. Our observations are consistent with previous studies reporting impaired spatial memory in PS19 males, but not females^76,77^, although one study observed impairments primarily in females at more advanced ages^75^.

The generally more severe spatial learning and neural activity deficits in males could arise from at least two non–mutually exclusive mechanisms. First, local tau burden within the MEC—particularly in pyramidal neurons—rather than the overall extent of tau spread, may be a primary determinant of spatial learning impairment. This interpretation aligns with human imaging and neuropathological studies showing that local tau burden in the entorhinal-hippocampal circuit correlates with cognitive decline more strongly than measures of tau spread alone^159,160^.

Second, because sex differences in tau distribution across the MEC were modest, additional sex-dependent molecular and cellular responses to tau pathology likely contribute to the observed behavioral and neural activity differences. Previous studies showed that although PS19 males and females exhibit similar levels of pathogenic tau isoforms across the brain, males display higher tau phosphorylation, increased astrocyte activation, elevated inflammatory signaling, and more pronounced microglial transcriptomic alterations^74,75^. Additionally, hormonal factors may further modulate vulnerability. Estrogen exerts neuroprotective effects in AD models^161^ and in mouse models of other conditions, such as stress^162^ and chronic liver disease^163^, consistent with the increased risk of post-menopausal women developing AD^164,165^ and with the relative preservation of spatial learning in female PS19 mice. Human transcriptomic studies also indicate that females exhibit distinct gene expression programs associated with tau pathology^166^, which may confer greater resilience to tau-related dysfunction.

Together, our findings highlight sex as a critical biological variable shaping tau-associated circuit dysfunction and spatial memory impairment, with implications for understanding sex-specific vulnerability in AD models^167,168^ and, ultimately, in human AD^169^.

### Clinical implications of our study

Our work supports a model in which tau pathology disrupts EC circuit function during aging and early in AD, leading to impairments in spatial cognition. By establishing a mechanistic link between tau burden, EC neural dynamics, and learning behavior, our study positions EC activity as both a clinically relevant biomarker for the diagnosis of early AD and a potential therapeutic target for preserving cognitive function in aging and AD.

First, our study identifies EC neural activity as a promising functional biomarker for aging-and early AD-related tau pathology. We show that EC activity is disrupted at a network level early in disease progression, closely correlates with local pTau burden, robustly predicts spatial learning performance, and reliably distinguishes tauopathy mice from controls—all prior to overt neurodegeneration. These results are highly relevant to human aging and disease, as converging clinical studies have reported impaired grid-cell-like representations in individuals at risk for AD^9,11^, as well as abnormal functional MRI activity of medial temporal lobe (MTL), including EC, associated with elevated tau burden and age-related memory decline^170^. Moreover, the EC activity abnormalities identified here align with deficits in path integration, which is a behavioral marker correlated with tau pathology in healthy adults^27^ and sensitively predicts AD risk before the onset of clinically detectable cognitive impairment^136,171,172^. Together, these findings indicate that EC functional measures may provide an early readout of tau pathology progression that complements established molecular and structural biomarkers, such as tau PET imaging and MTL atrophy^173,174^.

Second, our results suggest that EC circuit dysfunction itself represents a clinically actionable target for the treatment of AD-related tauopathy. Beyond molecular pathology, we identify specific neural activity features—including impaired speed modulation, unstable spatial representations, strong cue anchoring, reduced spatial selectivity, and altered activity levels—that are tightly linked to memory impairment. These findings raise the possibility that therapeutic strategies aimed at restoring EC network function—such as neuromodulation^174^, targeted stimulation^175,176^, or structured behavioral interventions^177,178^—could mitigate cognitive decline even when tau pathology is already present. Such circuit-level approaches could therefore enhance or extend the efficacy of disease-modifying therapies by stabilizing neural computations underlying spatial memory. Thus, our study will motivate future strategies to preserve or enhance EC function.

## ACKNOWLDEGMENTS

We thank all colleagues in the Gu lab for supporting the work, Dr. Michiyo Iba and Dr. Eliezer Masliah for their advice regarding the PS19 mouse model, Dr. Paul Taylor, Dr. Ben Engelhard, and Dr. Francisco Pereira for their help with the general linear modeling of animal behavior, and Dr. Huaibin Cai, Dr. Yun Li, Dr. Michiyo Iba, and Dr. Eliezer Masliah for constructive suggestions on the manuscript. This research was supported by the Intramural Research Program of the National Institutes of Health (NIH) to YG (ZIA NS009415). The contributions of the NIH authors were made as part of their official duties as NIH federal employees, are in compliance with agency policy requirements, and are considered Works of the United States Government. However, the findings and conclusions presented in this paper are those of the authors and do not necessarily reflect the views of the NIH or the U.S. Department of Health and Human Services.

## AUTHOR CONTRIBUTIONS

Conceptualization: TJM, YG

Methodology: TJM, KC, YG

Investigation: TJM, KC, JT, GW

Formal Analysis: TJM, KC, LC

Visualization: TJM, KC

Writing – original draft: TJM, YG

Writing – review & editing: TJM, YG

Funding acquisition: YG

Project administration: YG Supervision: YG

## DATA AVAILABILITY

All data used for figures and analysis in the study are available upon request from the lead contact (Dr. Yi Gu).

## CODE AVAILABILITY

Custom MATLAB scripts used for data analysis and figures are available on GitHub (https://github.com/tjmalone/Malone2026).

## DECLARATION OF GENERATIVE AI AND AI-ASSISTED TECHNOLOGIES IN THE WRITING PROCESS

During the preparation of this work the authors used ChatGPT 5.2 Pro in order to assist in shortening the manuscript. After using this tool, the authors reviewed and edited the content as needed and take full responsibility for the content of the published article.

## Supplementary Information

**Figure S1:**
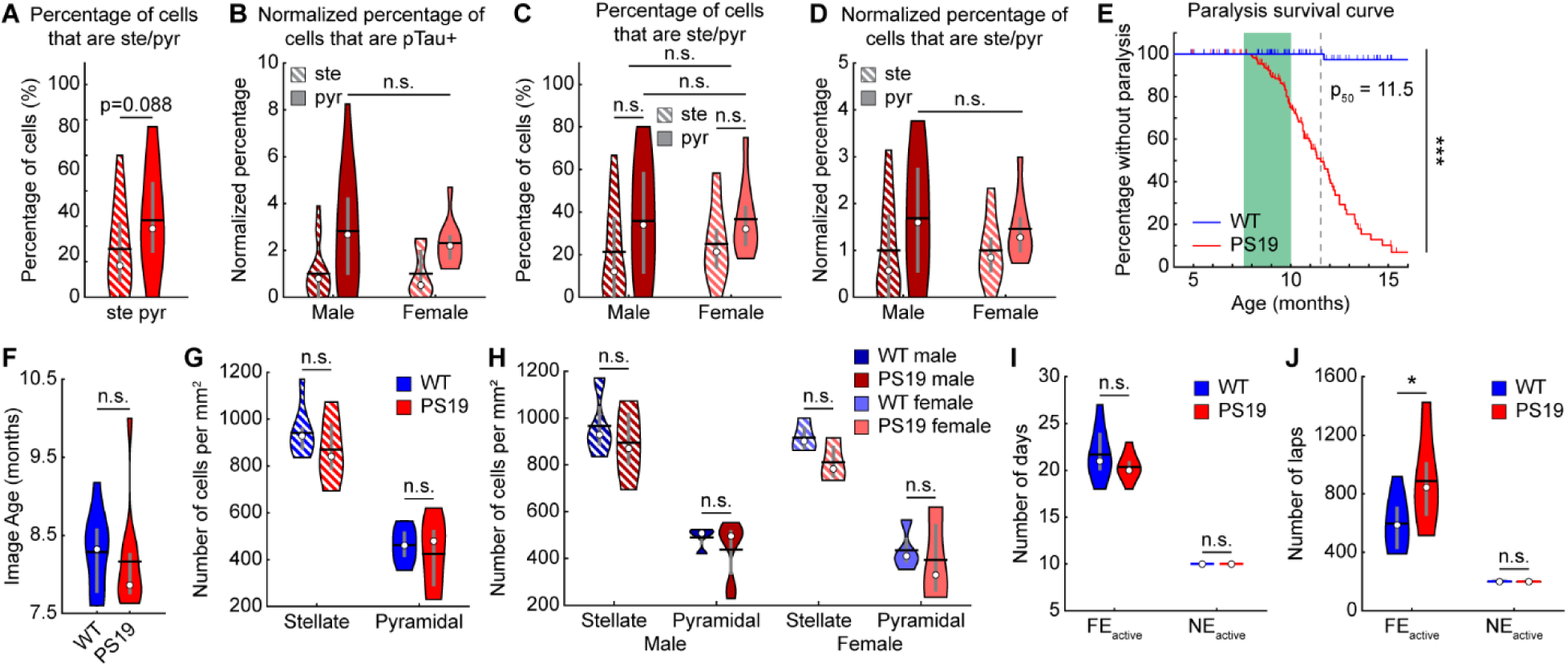
Pyramidal cells in PS19 mice show increased tau pathology. Related to Figure 1. **A.** Percentage of pTau+ cells overlapping with stellate (ste) and pyramidal (pyr) cells in all PS19 mice. **B.** Percentage of stellate and pyramidal cells overlapping with pTau+ cells in PS19 mice split by sex and normalized by stellate level for each sex. **C, D.** Percentage of pTau+ cells overlapping with stellate and pyramidal cells in PS19 mice split by sex. **C.** Raw overlap percentage. **D.** Normalized by stellate level for each sex. **A-D.** Paired/Unpaired Student’s t-test. **E.** Survival curve representing cumulative onset of hindlimb paralysis in WT and PS19 mice. Dashed line represents median PS19 value (p_50_). Green box represents age range of imaged mice. Ticks represent censored data points. Log-rank test. **F.** Mouse age at the start of imaging. **G, H.** Number of stellate and pyramidal cells per area in all mice (**G**) or for each sex (**H**). **I, J.** Number of training or imaging days (**I**) or laps (**J**) experienced in the familiar enviorment during active learning (training and imaging) or in the novel enviornment during imaging. **F-J.** Unparied Student’s t-test. \**p* ≤ 0.05, \*\**p* ≤ 0.01, \*\*\**p* ≤ 0.001. In violin plots, horizontal line represents mean, white circle represents median, and whiskers represent interquartile range. Statistical details can be found in **Table S1**.

**Figure S2:**
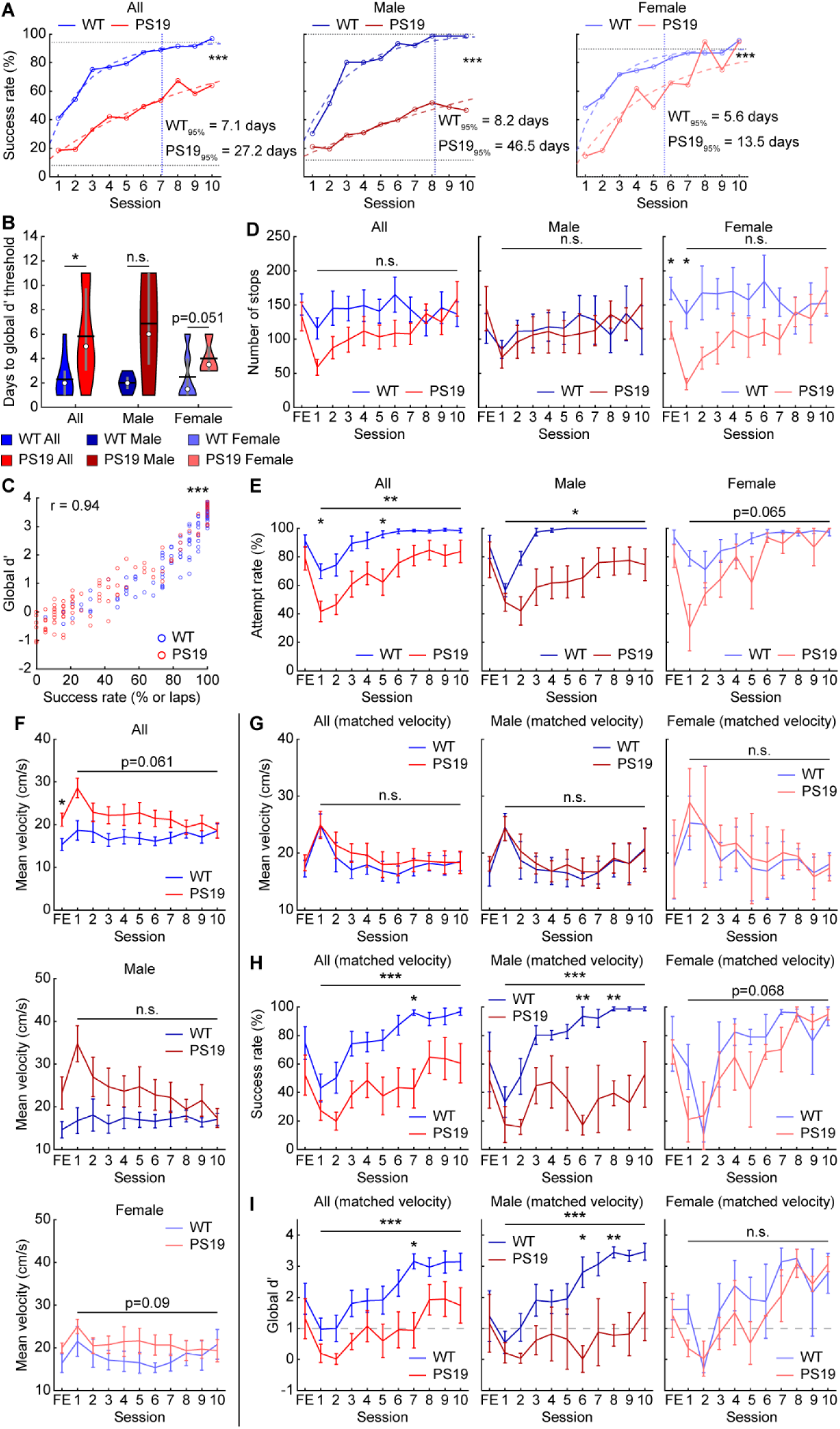
Male PS19 mice show behavioral deficits in an active spatial learning task. Related to Figure 1. **A.** Percentage of laps that mice successfully stopped to receive water reward across learning in all mice (left), male mice (middle), and female mice (right). WT_95%_ and PS19_95%_: number of days to reach 95% of plateau value. Solid lines and circles: mean success rate of each group from Fig. 1N. Dashed colored curves: fit of one-phase association exponential. Horizontal dotted lines: mutual baseline and plateau values used for exponential fit. Vertical dotted line: number of days for WT mice to reach 95% of plateau value. Extra sum-of-squares F test. All mice: WT*_K_* = 0.42, PS19*_K_* = 0.11. Male mice: WT*_K_* = 0.37, PS19*_K_* = 0.064. Female mice: WT*_K_* = 0.53, PS19*_K_* = 0.22. **B.** Number of days for global reward zone discrimination (global d’) to reach the learning threshold of 1. Unpaired t-test. **C.** Correlation between success rate and global d’ for all mice and sessions. Pearson correlation was calculated for both genotypes combined. **D.** Number of stops per session across learning in all mice (left), male mice (middle), and female mice (right). **E.** Percentage of laps that mice attempted to receive water reward across learning in all mice (left), male mice (middle), and female mice (right). An attempt was defined as 0.5 seconds (decreased from 1s) with speed below 5 cm/s (increased from 1 cm/s) within a 100 cm zone centered on the true 50 cm reward zone. **F.** Mean session velocity across learning in all mice (top), male mice (middle), and female mice (bottom). **G-I.** Mean session velocity (**G**), success rate (**H**), and global d’ (**I**) across learning in velocity matched mice for all mice (left), male mice (middle), and female mice (right). Velocity was matched by selecting pairs of mice on each day with similar velocity (within 5 cm/s). **D-I.** Horizontal gray bars indicate p values for the group difference (**D-F:** two-way repeated measures ANOVA, **G-I:** ordinary two-way ANOVA). Individual time points are compared using Student’s t-test with Bonferroni-Holm correction. \**p* ≤ 0.05, \*\**p* ≤ 0.01, \*\*\**p* ≤ 0.001. In violin plots, horizontal line represents mean, white circle represents median, and whiskers represent interquartile range. Error bars in line plots represent mean ± sem. Statistical details can be found in **Table S1**.

**Figure S3:**
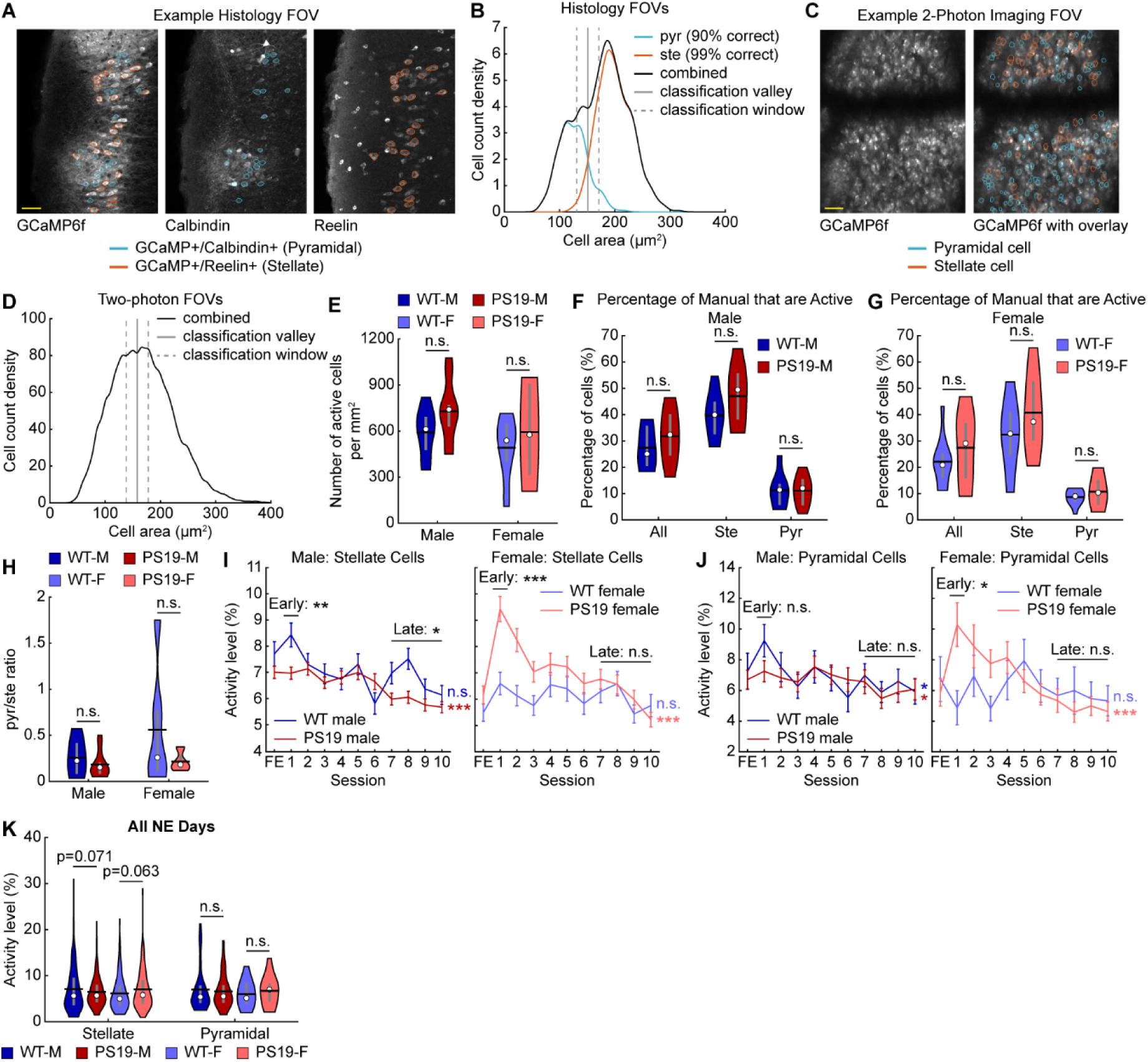
MEC cells of PS19 mice show hyper- or hypoactivity depending on molecular identity and sex. Related to Figure 2. **A.** Example histology FOV showing the overlap of reelin (stellate cell marker) and calbindin (pyramidal cell marker) with GCaMP6f. Cells were manually outlined if they had both GCaMP6f and reelin or calbindin staining. Used to validate differentiation of stellate and pyramidal cells based on cell area. Scale bar: 50µm **B.** Histology validation using area to differentiate stellate (ste) and pyramidal (pyr) cells. Colored lines indicate the true size distribution of stellate and pyramidal cells. The true positve rate of stellate and pyramidal cell idenification segementation (gray lines) using the combined area distribution (black line) is indicated. **C.** Example calcium imaging FOV with no overlay (left) and with an overlay of manually outlined cells identified as pyramidal or stellate cells (right). Scale bar: 50µm **D.** The size distribution of manually outlined cells from the FE day of calcium imaging with the segmentation lines marked as in **B** (158 +/- 20µm). **E.** The number of active cells per area on the FE day split by sex. WT-M: WT Male, PS19-M: PS19 Male, WT-F: WT Female, PS19-F: PS19 Female **F, G.** The percentage of all manually outlined cells and manually outlined stellate and pyramidal cells in male (**F**) or female (**G**) mice that are active on the FE day. **H.** The ratio between the fraction of active cells that are identified as pyramidal cells and as stellate cells split by sex. A low value indicates stellate cell enrichment. **I, J.** Activity level across learning in male (left) and female (right) mice in stellate (**I**) and pyramidal cells (**J**). **K.** Activity level averaged across all NE days split by sex and morphological cell type. \**p* ≤ 0.05, \*\**p* ≤ 0.01, \*\*\**p* ≤ 0.001. In violin plots, horizontal line represents mean, white circle represents median, and whiskers represent interquartile range. Violin plots use unpaired Student’s t-test. Error bars in line plots represent mean ± sem. In line plots, horizontal gray bars indicate p values for the group difference (early learning: unpaired Student’s t-test, late learning: two-way repeated measures ANOVA), and p-values to the right indicate significant Pearson correlation of the mean value with respect to time. Statistical details can be found in **Table S1**.

**Figure S4:**
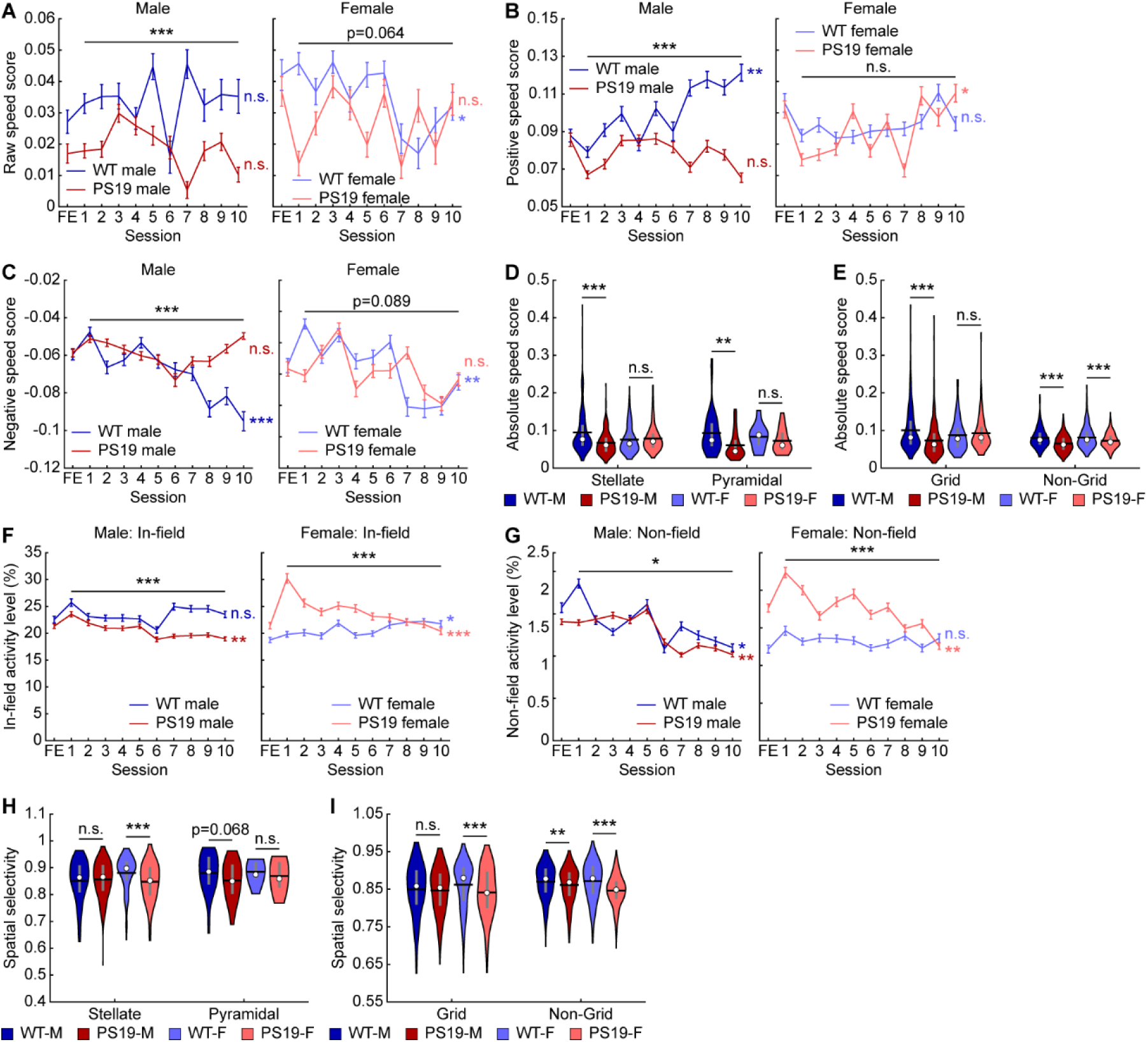
PS19 mice have altered speed modulation and spatial selectivity. Related to Figure 2. **A.** Raw speed score across learning in male (left) and female (right) mice. **B, C.** Positive (**B**) and negative (**C**) speed scores across learning in male (left) and female (right) mice. **D, E.** Absolute speed score averaged across all NE days split by sex and morphological (**D**) and functional (**E**) cell type. **F, G.** Activity level across learning in male (left) and female (right) mice within track regions with fields (In-field, **F**) and in other track locations (Non-field, **G**). **H, I.** Spatial selectivity averaged across all NE days split by sex and morphological (**H**) and functional (**I**) cell type. \**p* ≤ 0.05, \*\**p* ≤ 0.01, \*\*\**p* ≤ 0.001. In violin plots, horizontal line represents mean, white circle represents median, and whiskers represent interquartile range. Violin plots use unpaired Student’s t-test. Error bars in line plots represent mean ± sem. In line plots, horizontal gray bars indicate p values for the group difference (early learning: unpaired Student’s t-test, all learning days: linear mixed-effects model), and p-values to the right indicate significant Pearson correlation of the mean value with respect to time. Statistical details can be found in **Table S1**.

**Figure S5:**
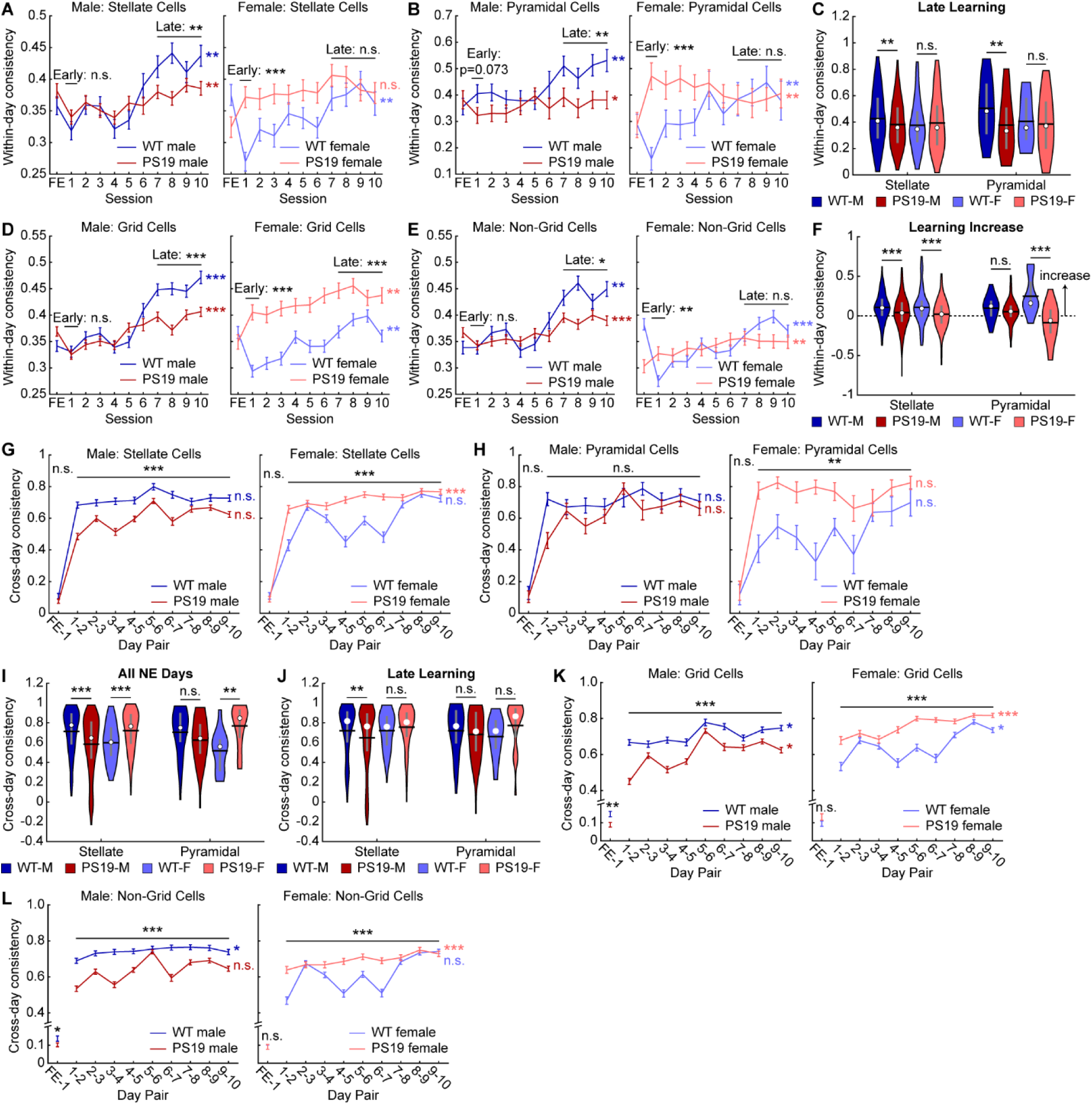
Male PS19 mice have impaired MEC within-day and cross-day activity consistency. Related to Figure 3. **A, B.** Within-day activity consistency across learning in male (left) and female (right) mice in stellate (**A**) and pyramidal cells (**B**). **C.** Within-day activity consistency averaged across late learning (days 7-10) split by sex and morphological cell type. WT-M: WT Male, PS19-M: PS19 Male, WT-F: WT Female, PS19-F: PS19 Female **D, E.** Within-day activity consistency across learning in male (left) and female (right) mice in grid (**D**) and non-grid cells (**E**). **F.** The increase in within-day activity consistency between early learning (day 1) and late learning split by sex and morphological cell type. **G, H.** Cross-day activity consistency across learning in male (left) and female (right) mice in stellate (**G**) and pyramidal cells (**H**). **I, J.** Cross-day activity consistency averaged across all learning days (**I**) and late learning (**J**) split by sex and morphological cell type. **K, L.** Cross-day activity consistency across learning in male (left) and female (right) mice in grid (**K**) and non-grid cells (**L**). \**p* ≤ 0.05, \*\**p* ≤ 0.01, \*\*\**p* ≤ 0.001. In violin plots, horizontal line represents mean, white circle represents median, and whiskers represent interquartile range. Violin plots use unpaired Student’s t-test. Error bars in line plots represent mean ± sem. In line plots, horizontal gray bars indicate p values for the group difference (early learning/ FE-1: unpaired Student’s t-test, late learning/all learning days: two-way repeated measures ANOVA or linear mixed-effects model), and p-values to the right indicate significant Pearson correlation of the mean value with respect to time. Statistical details can be found in **Table S1**.

**Figure S6:**
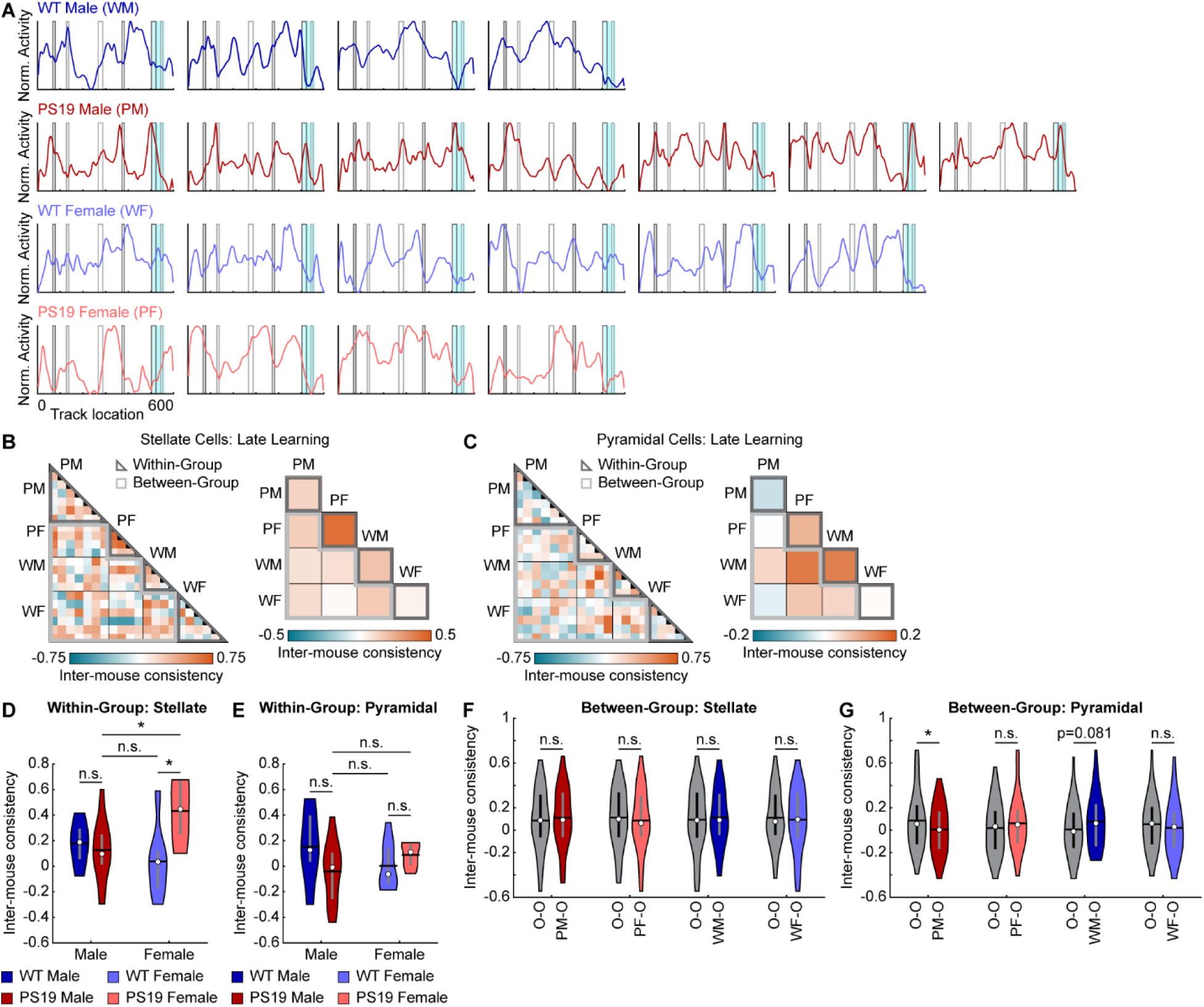
Male PS19 mice do not develop a cognitive map shared by other mouse groups. Related to Figure 4. **A.** Late-learning spatial activity maps as calculated in Fig. 4A for individual mice. From top to bottom: WT male, PS19 male, WT female, PS19 female. Black and grey rectangles represent left and right cues, respectively. Blue rectangle represents reward zone. **B, C.** Inter-mouse consistency for stellate (**B**) and pyramidal cells (**C**). Left: Pairwise inter-mouse consistency sorted by genotype/sex group. Right: Average inter-mouse consistency for each group-to-group block. **D, E.** Pooled inter-mouse consistency for pairwise within-group comparisons for stellate (**D**) and pyramidal (**E**) cells. Unpaired Student’s t-test with Bonferroni-Holm correction. **F, G.** Pooled inter-mouse consistency for pairwise between-group comparisons for stellate (**F**) and pyramidal (**G**) cells with each genotype/sex group serving as the reference group compared to the other genotype/sex groups (O) as indicated. Unpaired Student’s t-test. \**p* ≤ 0.05, \*\**p* ≤ 0.01, \*\*\**p* ≤ 0.001. In violin plots, horizontal line represents mean, white circle represents median, and whiskers represent interquartile range. Statistical details can be found in **Table S1**.

**Figure S7:**
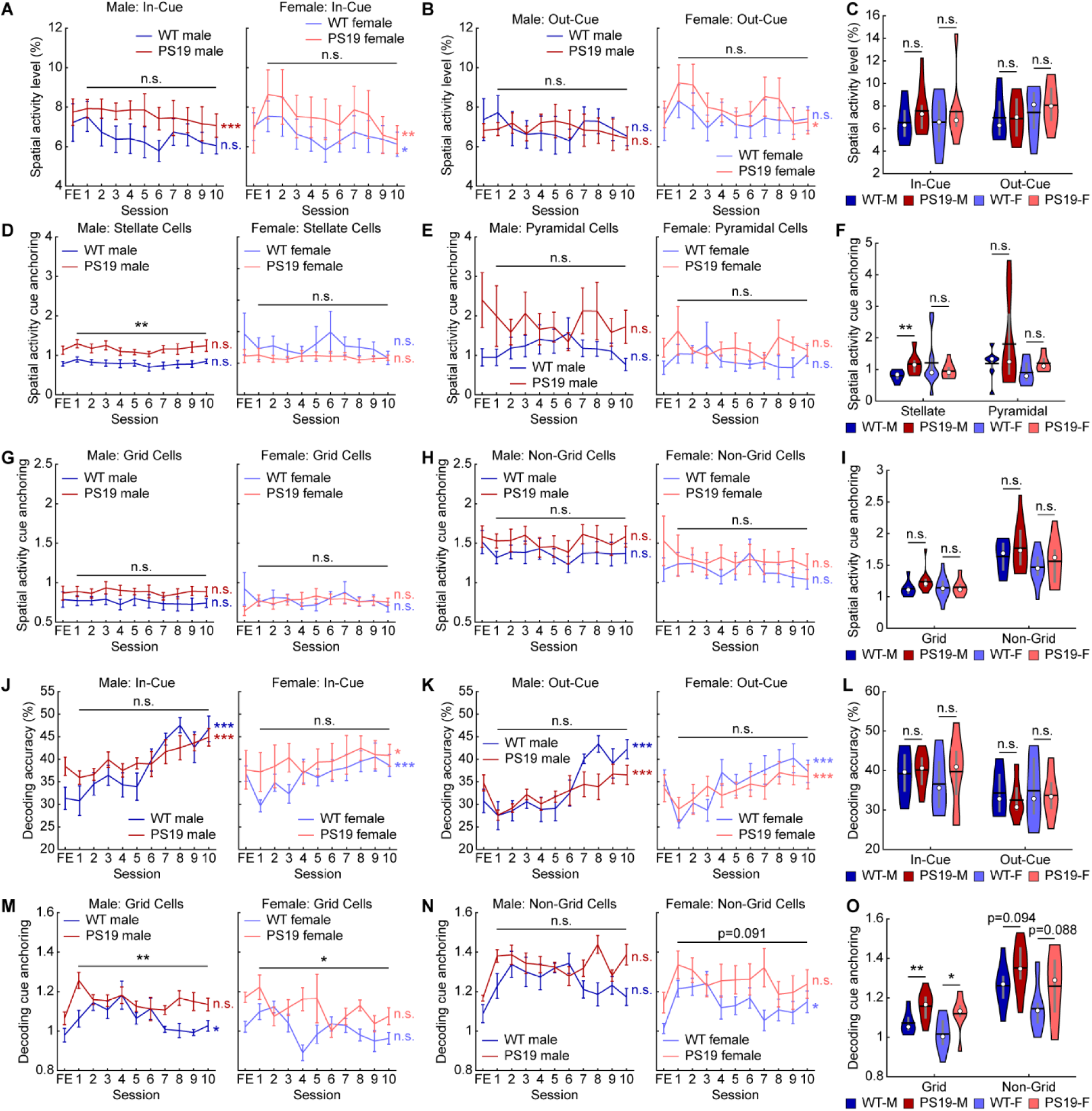
Spatial activity of male PS19 mice is anchored to visual cues. Related to Figure 5. **A, B.** Spatial activity level within in-cue (**A**) and out-cue (**B**) regions of the track across learning in male (left) and female (right) mice. **C.** Spatial activity level within in-cue and out-cue regions of the track averaged across all learning days. WT-M: WT Male, PS19-M: PS19 Male, WT-F: WT Female, PS19-F: PS19 Female **D, E, G, H.** Spatial activity cue anchoring across learning in male (left) and female (right) mice in stellate (**D**), pyramidal (**E**), grid (**G**), and non-grid (**H**) cells. **F, I.** Spatial activity cue anchoring averaged across all learning days split by morphological (**F**) and functional (**I**) cell type. **J, K.** Decoding accuracy within in-cue (**J**) and out-cue (**K**) regions of the track across learning in male (left) and female (right) mice. **L.** Decoding accuracy within in-cue and out-cue regions of the track averaged across all learning days. **M, N.** Decoding cue anchoring across learning in male (left) and female (right) mice in grid (**M**) and non-grid (**N**) cells. **O.** Decoding cue anchoring averaged across all learning days split by functional cell type. \**p* ≤ 0.05, \*\**p* ≤ 0.01, \*\*\**p* ≤ 0.001. In violin plots, horizontal line represents mean, white circle represents median, and whiskers represent interquartile range. Violin plots use unpaired Student’s t-test. Error bars in line plots represent mean ± sem. In line plots, horizontal gray bars indicate p values for the group difference (linear mixed-effects model), and p-values to the right indicate significant Pearson correlation of the mean value with respect to time. Statistical details can be found in **Table S1**.

**Figure S8:**
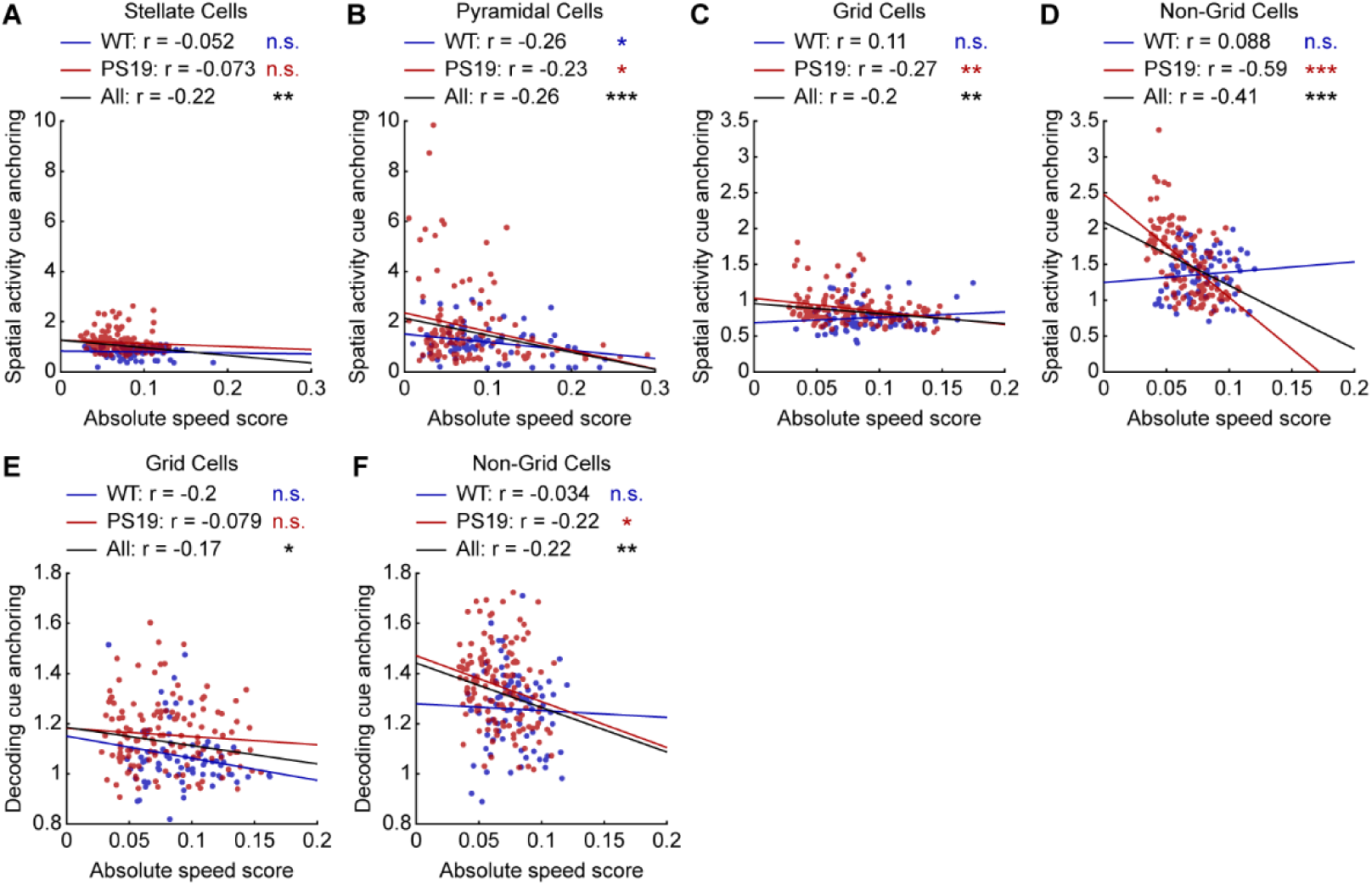
Speed modulation is correlated with cue anchoring in male PS19 mice. Related to Figure 5. **A-F.** Correlation between absolute speed score and spatial activity cue anchoring (**A-D**) or decoding cue anchoring (**E, F**) in stellate (**A**), pyramidal (**B**), grid (**C, E**), and non-grid (**D, F**) cells in male mice. Each point represents the activity from one FOV from one session. Pearson correlation. \**p* ≤ 0.05, \*\**p* ≤ 0.01, \*\*\**p* ≤ 0.001. Statistical details can be found in **Table S1**.

**Figure S9:**
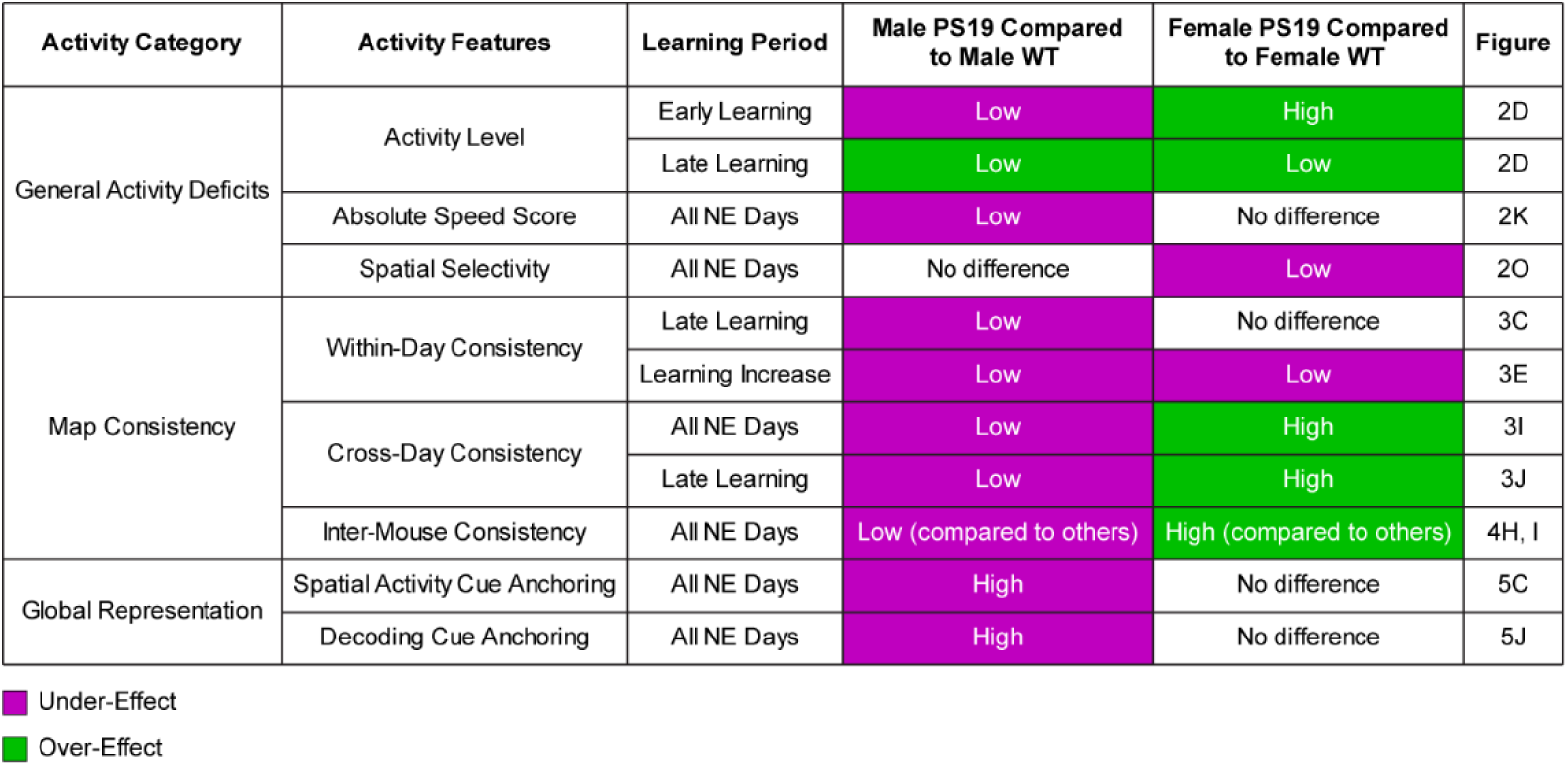
Summary table of activity under-effects and over-effects in PS19 mice relative to sex-matched WT mice.

**Figure S10:**
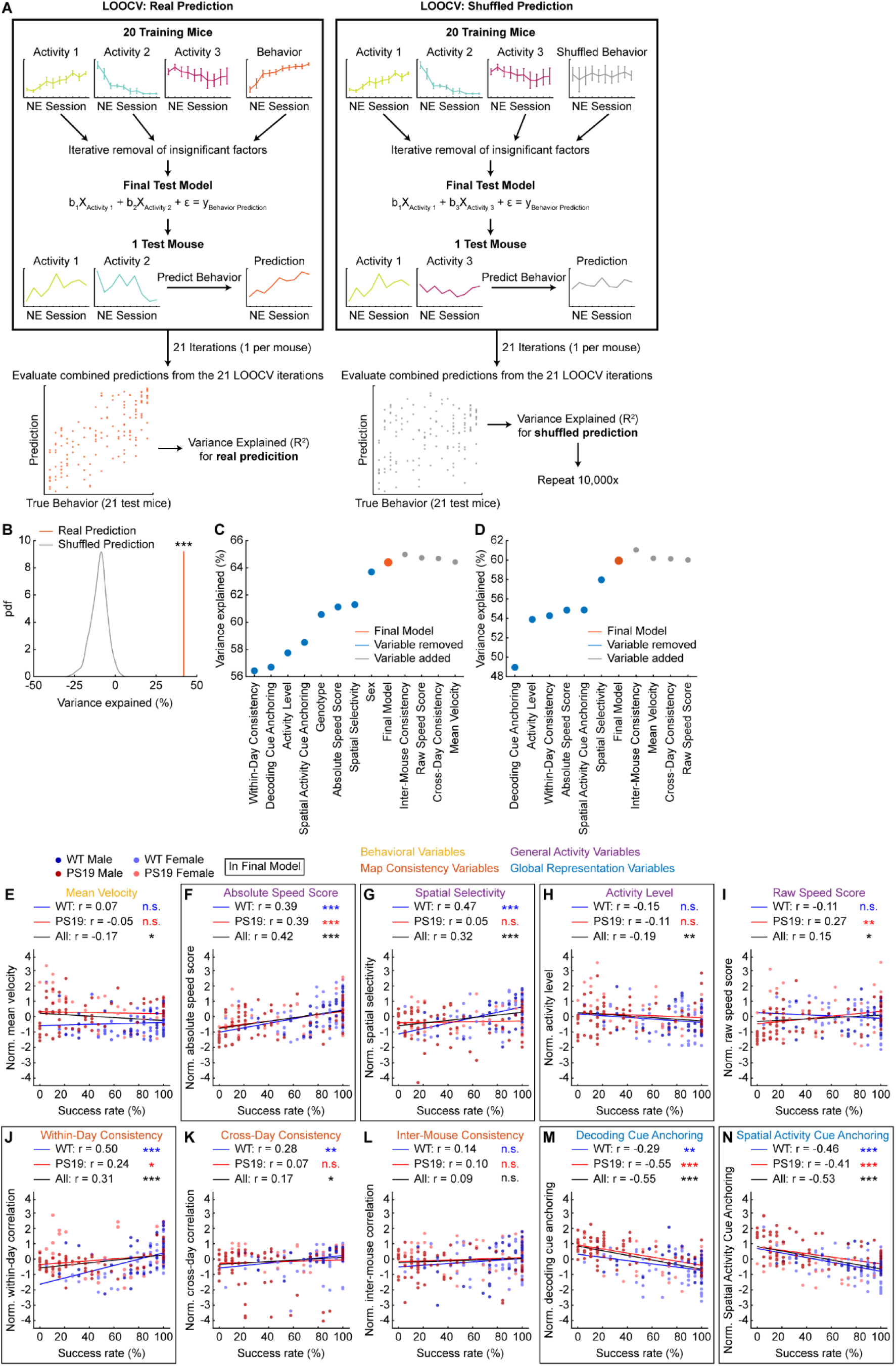
GLM model selection to predict mouse behavior using MEC activity. Related to Figure 7. **A.** Schematic of final model selection method. **B.** Evaluation of model selection method by comparing model predictions based on true behavior with shuffled behavior models. **C.** Effect of removing variables from final model or adding excluded variables back to final model on percentage of behavioral variance explained (R^2^). **D.** Similar to **C** but excluding sex and genotype from all models. **E-N.** Correlation between normalized predictor variables and success rate. Each point represents one session from one mouse. Dot colors indicate sex and genotype. Final model variables are outlined in black. Pearson correlations were calculated for all mice, all WT mice, and all PS19 mice. **E**: Mean velocity. **F**: Absolute speed score **G**: Spatial selectivity. **H**: Activity level. **I**: Raw speed score. **J**: Within-day consistency. **K**: Cross-day consistency. **L**: Inter-mouse consistency. **M**: Decoding cue anchoring. **N**: Spatial activity cue anchoring. \**p* ≤ 0.05, \*\**p* ≤ 0.01, \*\*\**p* ≤ 0.001. Statistical details can be found in **Table S1**.

**Figure S11:**
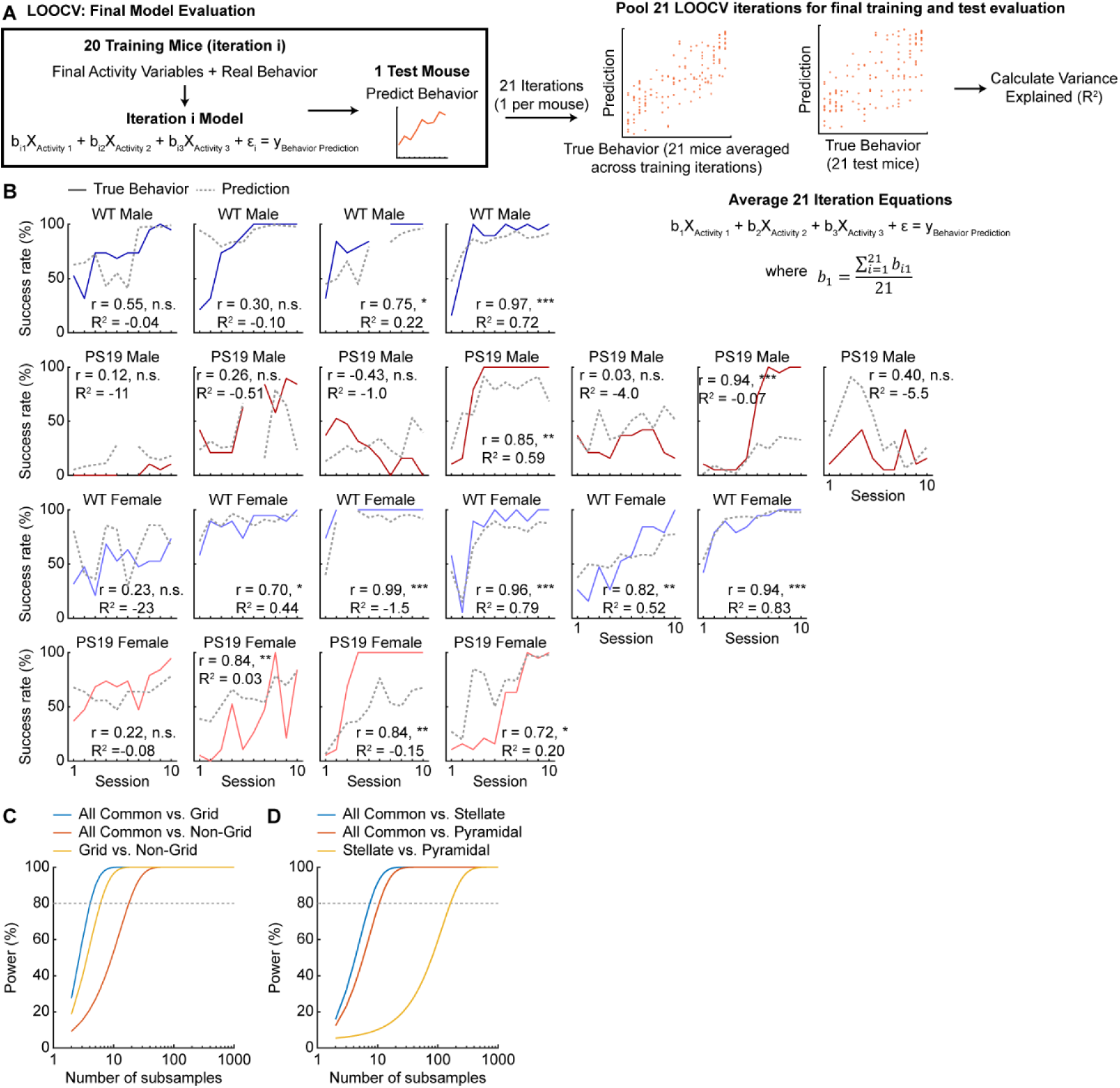
MEC activity changes are predictive of behavioral deficits in PS19 mice. Related to Figure 7. **A.** Schematic of final model evaluation method. **B.** Comparison of mean success rate (colored lines) and model predictions (dashed grey lines) across learning for individual mice (from top to bottom: WT male, PS19 male, WT female, PS19 female). Test predictions from single LOOCV iterations using the left-out mouse were used to calculate prediction. **C.** Power analysis comparing the distributions of behavioral variance explained of the subsampled all-common-cell, grid cell, and non-grid cell activity data to determine the probability of obtaining a significant result if a true difference exists for different numbers of subsamples. Dashed line corresponds to 80% power (common vs. grid = 5 subsamples, common vs. non-grid = 18 subsamples, grid vs. non-grid = 7 subsamples). **D.** Similar to **C**, but for all-common-cell, stellate cell, and pyramidal cell activity data (common vs. stellate = 8 subsamples, common vs. pyramidal = 11 subsamples, stellate vs. pyramidal = 162 subsamples). \**p* ≤ 0.05, \*\**p* ≤ 0.01, \*\*\**p* ≤ 0.001. Statistical details can be found in **Table S1**.

### METHODS

#### Animals

All animal procedures were performed in accordance with animal protocol 1524 approved by the Institutional Animal Care and Use Committee (IACUC) at NIH/NINDS. PS19 mice^58^ (B6;C3-Tg(Prnp-MAPT*P301S)PS19Vle/J, JAX stock #008169) were crossed with GP5.3 mice^67^ (C57BL/6J-Tg(Thy1-GCaMP6f)GP5.3Dkim/J, JAX stock #028280). The F1 generation of this cross was used for all experiments. Mice positive for GCaMP and human P301S will be referred to as PS19 and mice positive for GCaMP but negative for human P301S will be referred to as wild-type (WT). The two-photon imaging experiments for environment learning used 11 PS19 mice (7 males and 4 females) and 10 WT mice (4 males and 6 females). These imaging mice ranged from 7.6-10.0 months old at the start of imaging. Mice for immunohistochemistry experiments included a subset of the imaging mice and additional mice. The additional histology mice did not necessarily express GCaMP. Detailed sex and age information can be found below for each histology analysis. Mice were maintained on a reverse 12-hr on/off light schedule with all experiments being performed in the light off period. Animals were house at a temperature of 70–74°F and 40–65% humidity.

##### Paralysis Onset

The age at which paralysis onset first occurred as determined by onsite veterinary staff was recorded for all F1 mice in the colony, after which mice were humanely euthanized. The age of death was recorded for all mice 5 months or older regardless of the cause of death. Living mice were also included in the analysis. Ages for deaths other than after hindlimb paralysis and for living mice were used as censored events. In total, 96 WT mice (44 females and 52 males) and 208 PS19 mice (103 females and 105 males) were included in this analysis.

#### Rodent Surgeries

##### Microprism Construction

Microprism construction procedures were similar to those described previously^68,179^. A canula (MicroGroup, 304H11XX) was attached to a circular cover glass (3mm, Warner Instruments, 64-0720). A right angle microprism coated with aluminum on the hypotenuse (1.5mm, OptoSigma, RBP3-1.5-8-550) was then attached to the opposite cover glass side. All attachments were performed using UV-curing optical adhesive (ThorLabs, NOA81).

##### Microprism Implantation Surgery

Microprism implantation procedures were similar to those described previously^68,179^. Mice were anesthetized using a tabletop laboratory animal anesthesia system (induction: 3% isoflurane, 1 L/min oxygen, maintenance: 0.5–1.5% isoflurane, 0.7 L/ min oxygen, VetEquip, 901806) and surgery was performed on a stereotaxic alignment system (Kopf Instruments, 1900). A homeothermic pad and monitoring system (Harvard Apparatus, 50-7220 F) was used to maintain a body temperature of 37 °C. After anesthesia induction, dexamethasone (2 mg/kg, VetOne, 13985-533-03) and saline (500 μL, 0.9% NaCl, McKesson, 0409-4888-50) were administered by intraperitoneal (IP) injection, and slow-release buprenorphine (1 mg/kg, ZooPharm, Buprenorphine SR-LAB) was administered subcutaneously. Enroflox 100 (10 mg/mL, VetOne, 13985-948-10) was used as an anti-microbial wash just after the skull was exposed and just prior to sealing the skull. All insertions were performed on the left hemisphere, aligning with previous observations of more favorable vasculature^179^. A 3 mm craniotomy was performed centered at 3.4 mm lateral to the midline and 0.75 mm posterior to the center of the transverse sinus (approximately 5.4 mm posterior to the bregma). A durotomy was then performed over the cerebellum. Mannitol (3 g/kg, Millipore Sigma, 63559) was administered by IP prior to the durotomy. The microprism was inserted into the transverse sinus and sealed to the skull with n-butyl cyanoacrylate tissue adhesive (Vetbond, 3 M, 1469SB). The exposed skull was also coated with Vetbond. A single-sided steel head plate for head fixation was mounted to the right side of the skull opposite the craniotomy. Finally, the prism and head plate were adhered to the skull with dental cement (Metabond, Parkell, S396).

#### Virtual Reality Setup

For all behavioral experiments, a customized virtual reality (VR) setup was used, which projects a one-dimensional (1D) virtual environment based on the running of a mouse, similar to that described previously^49,179^. Mice were head-fixed onto an air-supported polystyrene ball (8” diameter, Smoothfoam) using the mounted head plate. The ball rotated on an axle, allowing only forward and backward rotation. The virtual environment was projected onto a hemispherical dome filling the visual field of mice (270° projection). An optical flow sensor (Paialu, paiModule_10210) with infrared LEDs (DigiKey, 365-1056-ND) was used to measure the rotation of the ball and thereby control the motion of the virtual environment. The optical flow sensor output to an Arduino board (Newark, A000062), which transduced the motion signal to the computer controlling the VR. An approximately 4 μl water reward was provided via a lick tube using a solenoid. The solenoid was controlled using a Multifunction I/O DAQ (National Instruments, PCI-6229). The virtual environments were generated and projected using ViRMEn software (Princeton, version 2016-02-12)^180^. Imaging and behavior data were synchronized by recording a voltage signal of behavioral parameters from the VR system using the DAQ. ViRMEn environments were updated at 60 Hz. The DAQ input/output rate was 1 kHz. The synchronization voltage signal was updated at 20 kHz. Final behavioral outputs were matched to the imaging frame rate (15 Hz, see below) for synchronization.

Environments were colored blue and projected through a blue wratten filter (Kodak, 53-700) to reduce contamination of the imaging path with projected light. Virtual environments were 600 cm 1D linear tracks with patterned walls and patterned visual cues at fixed locations (**Fig. 1M**). At the end of the track, mice were immediately teleported to the start of the track.

#### Immunohistochemistry

Mice were anesthetized with a ketamine (200mg/kg, VetOne, 13985-584-10) and xylazine (20mg/kg, VetOne. 13985-612-50) cocktail and were transcardially perfused with 4% paraformaldehyde (PFA, Electron Microscopy Sciences,15713) in phosphate buffer solution (PBS, Corning, 46-013-CM). Brain tissues were dissected and fixed in 4% PFA in PBS overnight at 4°C. Sagittal slices (40µm thick) were prepared using a VT1200S vibratome (Leica Biosystems). Slices were washed in PBS (4x 10 minutes) and then blocked with 10% Bovine serum albumin (BSA, Millipore Sigma, A3294), 0.5% triton X-100 (Millipore Sigma, T9284), and PBS for 1 hour at room temperature. Primary and secondary antibodies were diluted in 10% BSA, 0.5% triton, and PBS. Slices were incubated in diluted primary antibodies for 48 hours at 4°C, then washed in PBS (4x 10 minutes). Slices that required anti-goat secondary antibodies were incubated in diluted anti-goat secondary antibodies (1:500 dilution) for 1.5-2 hours at room temperature, washed in PBS (3x 10 minutes), incubated in other secondary antibodies (1:500 dilution) for 1.5-2 hours at room temperature, and washed in PBS (3x 10 minutes). Slices that did not require anti-goat secondary antibodies were incubated in the diluted secondary antibodies (1:500 dilution) for 1.5-2 hours at room temperature, then washed in PBS (3x 10 minutes). Slices were mounted with mounting medium (Vector Laboratories, H-1000-10). Primary antibodies used include goat anti-reelin (1:400, R&D Systems, AF3820), rabbit anti-Calbindin (1:500, Abcam, AB108404), and mouse monoclonal IgG1 anti-Phospho-Tau (1:1000, ThermoFisher Scientific, MN1020). Secondary antibodies included Alexa 405 conjugated donkey anti-goat antibody (Invitrogen, A48259), 568 conjugated goat anti-rabbit antibody (Invitrogen, A-11036), and Alexa 647 conjugated goat anti-mouse IgG1 antibody (Invitrogen, A-21240).

##### Imaging

Histology slices were imaged using a Leica M165 FC microscope for the phospho-tau (pTau) accumulation with age quantification and using a Zeiss 880 spectral confocal for all other histology analyses. 3 to 4 image stacks were taken on the confocal microscope of the medial entorhinal cortex (MEC) along the dorsal-ventral axis and were stitched together using the “Pairwise stitching” plugin in ImageJ (Fiji, version 1.53q). 3-10 focal planes (Z-planes) were imaged for each image stack.

##### Quantification of pTau Accumulation with Age

6 WT Males, 21 PS19 males, 6 WT females, and 11 PS19 females ages 2 to 10 months were stained in 3 staining batches. 16 of these mice were in the 2-photon imaging experiments. 1 Z-plane of the image stack was analyzed per slice. The dorsal 1.5mm region of MEC layer 2 was selected manually and the intensity of pTau immunostaining of the MEC layer 2 region was measured in ImageJ. A cutoff for pTau+ pixels in layer 2 was calculated for each of the three staining batches. The 99^th^ percentile value of all the pixels in layer 2 was measured for each 2- and 3-month PS19 mouse in each of the staining batches. The cutoff for each batch was the average of these intensity values. The percentage area that was pTau+ was the number of pTau+ pixels above the intensity cutoff divided by the total number of pixels in the MEC layer 2 region. The pTau intensity in the pTau+ region was the average pixel value in MEC layer 2 after subtracting the cutoff intensity and excluding negative pixel values.

##### Quantification of pTau Overlap with Reelin and Calbindin Cells

9 PS19 males and 5 PS19 females ages 8 to 10 months were stained in 4 staining batches. 10 of these mice were in the 2-photon imaging experiments. 2 Z-planes of the image stack were analyzed per slice. The dorsal 1.5mm region of MEC layer 2 and reelin+, calbindin+, and pTau+ cells inside this region were selected manually in ImageJ. Reelin and calbindin selections were overlaid with pTau+ cell selections. Reelin and calbindin selections were manually determined to be pTau+ by counting overlapping regions of interest (ROIs). pTau+ cells that overlapped with both reelin and calbindin selections were excluded when calculating the percentages of reelin+ and calbindin+ cells that are pTau+ and pTau intensity. The pTau intensity of pTau+ cells that overlapped with reelin+ or calbindin+ cells was calculated as the average pixel value after background subtraction within each Z-plane.

##### Neurodegeneration Quantification

4 WT Males, 7 PS19 males, 4 WT females, and 3 PS19 females, ages 8 to 10 months were stained in 4 staining batches. 17 of these mice were in the 2-photon imaging experiments. 1 Z-plane of the image stack was analyzed per slice. The dorsal 1.5mm region of MEC layer 2 and reelin+ and calbindin+ cells inside this region were selected manually in ImageJ. Cells with less than 90% of their area inside the layer 2 ROI were removed. Neurodegeneration was quantified by counting the number of reelin+ and calbindin+ cells and dividing by the area of the layer 2 region to get the number of cells per area.

#### Behavior

##### Training

Water-restricted mice were first trained in a 1 m environment with a single reward at a fixed track location during acclimation to VR to encourage running. Once mice began to run (typically 50-100 laps/hour for two days), they were transitioned to a 6m environment with a single water reward at a fixed track location (**Fig. 1M**, passive FE). Mice were trained in the passive FE until they started to develop reward-predictive behavior as measured by predictive licking and predictive slowing (roughly 50% predictive licking and 60% predictive slowing, see “Predictive Licking” and “Predictive Slowing” for quantification), which has been shown previously to demonstrate familiarity with an environment^49,181^. Both WT and PS19 mice were able to reach these criteria. Mice then began training in an active task (active FE). The active FE was visually identical to the passive NE, but mice only received water reward if they stopped in an unmarked 50 cm reward zone centered at the reward location of the passive FE. A stop was defined as a velocity below 1 cm/s for at least 1 s. Mice could only receive one reward per lap. For the first 3-4 days, mice underwent manual training, consisting of manually stopping the mouse in the reward zone. When a mouse could stop on its own, it was further trained without manual stopping with a shortened stop period (0.5s). Once a mouse stopped on its own with the shortened stop period, the full 1s stop period was applied. Additional manual training was performed periodically as necessary to reinforce correct behavior.

Mice were trained in the active FE until they learned the active task (success rate >∼75%, see “Success/Attempt Rate” for quantification) and ran well in the active FE environment (>∼20 laps per hour). Because there was no a-priori assumption that PS19 mice would be able learn the active spatial task, mice (WT or PS19) that did not show learning of the active task were transitioned when they had been trained in the active task for a similar number of days to the average WT mouse (>∼17 days). Mice then underwent several days of baseline imaging in the active FE environment, until their behavior recovered to its pre-imaging level. Further behavioral analysis and analysis of neuronal activity used the FE imaging day with the highest success rate in order to ensure a baseline day representing neuronal activity in a well-learned environment. Mice were then imaged for 10 consecutive days in the novel environment (NE), which had a different arrangement of visual cues with different patterns and a 50cm reward zone in a different location (**Fig. 1M**). Mice were restricted to 20 laps per day in the NE. Because imaging could not be performed for the entirety of the first lap, the first lap was ignored for behavioral analysis as well.

##### Success/Attempt Rate

Success rate was measured as the percentage of laps that a mouse stopped in the reward zone. The success rate criteria used for analysis were identical to those used to determine when the mouse received reward (1 s with velocity less than 1 cm/s). To measure attempts to receive reward, we conducted a post-hoc analysis in which we relaxed the criteria for a successful stop in reward zone. Specifically, we simultaneously expanded the reward zone (from 50 cm to 100cm), increased the speed threshold defining a stop (from 1 cm/s to 5 cm/s), and reduced the amount of time required for a stop (from 1 s to 0.5 s).

##### Predictive Licking

Predictive licking (PL) was measured as the percentage of licks that occurred within a specific distance threshold (20 cm) prior to the reward delivery relative to all other locations (excluding 30 cm after reward). PL was only calculated for the passive FE to determine the transition to active learning.

##### Predictive Slowing

To calculate predictive slowing (PS), for each lap, the velocity changes between each data point were calculated along the track. The pre-reward acceleration value was calculated as the mean velocity changes (Δv) within a 90 cm window before the reward (for example 175 cm to 265 cm for a reward location at 265 cm). The rest of the track was analyzed using a rolling average of Δv for the same window size at 1 cm intervals, generating a series of comparison acceleration values. The slowdown percentile for a given lap was the percentile of the pre-reward acceleration within the comparison acceleration values from the rest of the track, such that a higher percentile signified a more negative acceleration (i.e., deceleration) and thus better PS. The rolling averages calculated for the comparison values excluded any window intersecting the edge of the track or areas close to the reward (from 90 cm before to 30 cm after reward delivery) to avoid edge effects and reward related behavior, respectively. Thus, the comparison windows ranged from 0–90 cm to 85–175 cm and 295-485 to 510-600 on the passive FE. The PS of a session was computed as the average slowdown percentile of all the laps within the session. PS was only calculated for the passive FE to determine the transition to active learning.

##### Global Discrimination Index

A global discrimination index (global d’) was calculated to measure the ability of a mouse to discriminate between the reward zone and other locations. The success rate was compared to the percentage of laps with stops in a rolling 50 cm window of track positions (the same size as the reward zone). For each comparison window an individual local d’ value was calculated as d’ = norminv(HR)-norminv(FA); where HR=Hit Rate or success rate in the reward zone, and FA=False Alarm Rate or the percentage of laps with a stop in a given comparison zone^85,86^, and norminv() is the inverse function of the cumulative Gaussian distribution. HR and FA were constrained between 1/2N and 1-1/2N, where N is the number of laps, because norminv is undefined for 0 or 1. Global d’ is the average d’ for all comparison windows. A threshold of d’ >1 was defined as the threshold for successful learning^85–87^. If mice never reached a global d’ of 1, the number of days to reach the learning threshold was considered to be 11 (total number of NE days + 1) as a penalty.

##### Mean Velocity

The mean velocity of a mouse for a given session was defined as the average velocity of all time points where the mouse was moving faster than 1 cm/s. This value was used for both analyzing mouse behavior and as a potential predictor variable for the general linear models predicting behavior.

##### Number of Stops per Session

The number of stops in a given session was calculated as the number of distinct instances when the velocity of a mouse went below 1 cm/s for at least 1 s. A given stop could be arbitrarily long and continued until the velocity rose above 1 cm/s again. Therefore, two stops could be close together in space and time but were required to have at least some period of higher velocity separating them.

##### Velocity Matching

To perform velocity matching, pairs of mice with similar velocity were matched on each day. For each sex group (all, male, female), the optimal set of cross-genotype mouse pairs was determined using the MATLAB (Mathworks, v2023b) matchpairs function, which finds the combination of pairs with the minimum total cost (velocity difference). All possible pairs of velocity difference between a WT and PS19 mice were initially calculated, but the cost of any pairs with a velocity difference greater than 5 cm/s was set to infinity. In addition, the cost of unpaired mice was set to 10^6^ (much higher than the velocity values). This method finds the maximum possible number of mouse pairs with a velocity difference less than 5 cm/s. The velocity-matched velocity, success rate, and global d’ were then calculated using only the mice that were part of a valid pair on each day.

##### Behavior Fitting

To calculate the rate of spatial learning, the mean success rate of each group was fitted with a one-phase association exponential function (Prism 9.3.1, GraphPad), where y is the success rate and x is day (NE day 1 to 10). For each sex group (all, male, female), we first determined if both WT and PS19 mice had the same baseline (*Y*_0_) and plateau (*P*) success rate using an extra sum-of-squares F test. This analysis found that for all sex groups, fitting the two genotypes with the same baseline and plateau values was preferred to fitting the genotypes separately (Extra sum-of-squares F test. All mice: *Y*_0_=8.0, *P*=94, p=0.63. Male mice: *Y*_0_=12, *P*=100, p=0.13. Female mice: *Y*_0_=0.0, *P*=90, p=1). We then re-fit each genotype with the baseline and plateau levels set at the previously determined mutual values and let K, the learning rate constant, vary. We again used an extra sum-of-squares F test to determine whether the genotypes were better fit with the same or different values of K. If the F test determined that the values of K are different, we could conclude that the genotype with the higher rate constant learned faster. Once the rate constant was derived, we calculated the number of days required to achieve 95% of the plateau (*T*_95%_) using the equation *T*_95%_ = ln(20/K). The *T*_95%_ directly corresponds to K but is more interpretable.

#### Two-Photon Imaging

Imaging was performed using an Ultima 2Pplus microscope (Bruker) configured with the above VR setup. A tunable laser (Coherent, Chameleon Discovery NX) set to a 920 nm excitation wavelength was used. Laser scanning was performed using a resonant-galvo scanner (Cambridge Technology, CRS8K). GCaMP fluorescence was isolated using a bandpass emission filter (525/25 nm) and detected using GaAsP photomultiplier tubes (Hamamatsu, H10770PB). A 16x water-immersion objective (Nikon, MRP07220) was used with ultrasound transmission gel (Sonigel, refractive index: 1.3359^182^; Mettler Electronics, 1844) as the immersion media.

The anterior-posterior (AP) and the medial-lateral (ML) angle of the prism (i.e., the angle of the surface of the prism along to the AP or ML direction of the mouse) relative to the head-fixed position of the mouse were measured prior to the first imaging session. The head plate holder and rotatable objective angles were set daily to align the objective with the prism in the AP and ML direction, respectively, such that the objective was parallel to the prism surface. Black rubber tubing was wrapped around the objective and imaging window to prevent light leakage into the objective.

Microscope control and image acquisition were performed using Prairie View software (Bruker, version 5.7). Raw data was converted to images using the Bruker Image-Block Ripping Utility. Imaging was performed simultaneously on two fields-of-view (FOV) using an electrotunable lens (ETL, Bruker). FOV were 750 μm x 750 μm and were separated by 35-43 μm in the z-plane. Each FOV was imaged at 15 Hz with 512 × 512 resolution. Average beam power at the front of the objective was typically 100–120mW. Imaging and behavior data were synchronized as described above. Only the activity on laps where imaging was acquired for the full lap was used for analysis. Due to technical issues, four mice were not successfully imaged for one NE day. These four sessions are included in the behavioral analysis but are excluded from the imaging and modeling analysis. To ensure comparable common cell identification (see “Cell Alignment”), an additional NE imaging day was used for the alignment of these mice, such that all mice were aligned with 11 imaging days.

#### Image Processing

Imaging data underwent a consecutive 2-frame rolling average and was processed as previously described using published MATLAB scripts^68^. Motion correction was performed using cross-correlation based, rigid motion correction. Identification of regions of interest (ROIs, active cells) with correlated fluorescence changes and extraction of the fluorescence time course of individual ROIs was performed using Suite2p (v0.11.1)^183^. The fractional change in fluorescence with respect to baseline (ΔF/F, referred to as activity level) was calculated as (F(t) – F0(t)) / F0(t)^179^. For each cell, significant calcium transients were identified using amplitude and duration thresholds, such that the false-positive rate of significant transient identification was 1%^184^. A final ΔF/F including only the significant calcium transients (as opposed to the raw ΔF/F) was used for further analysis, except as noted. A speed threshold was calculated by generating a 100-point histogram of all instantaneous velocities greater than 0 and taking the value twice the center of the first bin (approximately 1% of max positive speed). The ΔF/F points when the mouse was moving below the speed threshold were excluded from the following analyses. The mean ΔF/F (significant transients only, as mentioned above) for a cell was calculated as a function of position along the track in 5 cm bins.

##### Cell Alignment

All imaging sessions for a given mouse were aligned pairwise using an established probabilistic modeling method^185^ to identify common cell pairs between each pair of sessions. We first performed automatic pairwise rigid registration of the imaging sessions to identify common neuron pairs between pairs of imaging days. Manual registrations were applied to the imaging session pairs with failed automatic alignment, but common cell pairs were still identified using the same probabilistic modeling method. Pairwise registration results were combined to determine the common neurons across all 11 days of the spatial learning task. Due to the pairwise nature of the registration, a neuron could be aligned to different neurons on different days. Manual examination was used to select the optimal alignment results that display the highest level of anatomical footprint consistency across all imaging days and to exclude incorrect alignments.

#### Data Analysis

##### Stellate/Pyramidal Cell Classification

Cells were classified as stellate cells or pyramidal cells based on the bimodal distribution of their area similar to a previously described method^68^. Because previous studies used long-axis diameter of the cells instead of area, we first validated this method using histology slices of cells expressing GCaMP6f that were stained for reelin (a stellate cell marker) and calbindin (a pyramidal cell marker). Cells that expressed both GCaMP6f and either reelin or calbindin, but not both, were manually traced and their area was calculated. The valley between the peaks of the cell area was 151 μm^2^. To increase the confidence of cell classification, cells with areas smaller than 131 μm^2^ and larger than 171 μm^2^ (151 ± 20 μm^2^) were classified as pyramidal and stellate cells, respectively. We calculated the true positive rate of stellate and pyramidal cells as the number of cells identified as stellate or pyramidal cells, respectively, divided by the total number of cells identified as stellate or pyramidal cells (stellate cells: 99% true positive rate, pyramidal cells: 90% true positive rate).

For the imaging FOV, cells were manually traced from motion corrected maximal projections of each FOV on the FE imaging day. The valley between the two peaks of cell area was 158 μm^2^. To increase the confidence of cell classification, cells with areas smaller than 138 μm^2^ and larger than 178 μm^2^ (158 ± 20 μm^2^) were classified as pyramidal and stellate cells, respectively.

##### Matching Manually Identified Cells to Active Cells

To identify common cells as stellate or pyramidal cells, the ROIs of the active cells on the FE day were overlaid with the manually traced ROIs. An active cell and a manually drawn cell were determined to be the same cell if the centroid distance between the two ROIs was less than 4 μm and the area of the intersect of the ROIs divided by the area of the union of the ROIs (IoU) was less than 0.4. In the event that a given cell matched with multiple cells, the cell pair with the smallest centroid distance was used. All common cells that matched a manual cell on the FE day were assigned that identity across all days. Because not every active cell matched with a manual cell and not every manual cell was identified as stellate or pyramidal, the population of common cells identified as stellate or pyramidal cells is only a subset of the total common cell population.

##### Spatial Field Identification

To calculate significant spatial firing fields, regions of the track with significantly consistent activity, mean ΔF/F was compared to shuffles of the original ΔF/F as described previously^68^. Mean ΔF/F for a cell was spatially binned along the track in 5 cm bins and averaged across laps. A gaussian window of 3 spatial bins was applied to the averaged ΔF/F trace to smooth the data. Spatial fields were identified by comparing the amplitude of the original smoothed ΔF/F with a random ΔF/F distribution created by 1000 bootstrapped shuffled responses. Each bootstrapped shuffled response was generated by rotating the ΔF/F trace of the whole session so that for every time point of the recording, its track position was preserved but its calcium response was changed. The ΔF/F trace was rotated by starting the trace from random sample numbers chosen from the interval 0.05 × Nsamples to 0.95 × Nsamples, where Nsamples was the number of samples in the ΔF/F trace. A shuffled mean ΔF/F was calculated for each rotation. For each 5 cm bin, the pvalue equaled the percentage of shuffled mean ΔF/F that was greater than or equal to the real mean ΔF/F. Therefore, 1 - pvalue equaled the percentage of shuffled mean ΔF/F lower than the real mean ΔF/F. A spatial field was defined as a region of at least 3 consecutive 5 cm bins (except that the fields at the beginning and end of the track could have 2 bins) that had a mean ΔF/F higher than at least 80% of 1000 shuffles at the corresponding bins (1 - pvalue ≤ 0.2). Additionally, at least 20% of laps were required to have a significant calcium transient in the spatial field.

##### Cue Cell Classification

To identify cue cells, cue scores of each cell on a specific day session were calculated based on a previously published method^36^. The activity of the cell was first shifted to best match a cue template, which contained ones and zeros representing track areas with and without cues, respectively. A cue zone was identified for each cue by including the cue itself and the surrounding region expanded by half of the cue width on both sides. The correlations between shifted activity and cue template within individual cue zones were calculated and were further averaged as the cue score of the cell. 200 shuffled cue scores were generated for each cell using the same calculation but by randomizing the cue location in the cue template. Shuffled cue scores from all cells across all FOVs and imaging days for a given environment were pooled to generate an overall cue score distribution. A cell was identified as a cue cell if its cue score on the day was above the 90th percentile of the shuffle cue score distribution. While previous studies used a 95th percentile threshold^36,49,68^, a 90th percentile threshold was used here to better identify cells with cue-matched activity. Cue scores of cells were calculated toward left cues, right cues, and cues on both sides of the track. A cue cell could be identified based on any cue type. Cue cell identification was performed each day to exclude cue cells from the putative grid cell population (see “Grid Cell Classification”). Common cells were identified as true cue cells if they were identified as cue cells of any type (left, right, or both sides) on more than half of imaging days (≥ 6).

##### Grid Cell Classification

Grid cells were identified within each recording day based on the following criteria. (1) Grid cells need to have at least two spatial fields on the track. (2) Each grid cell must have more than L/(5w) transitions between in-field and out-of-field periods, where L represents track length and w represents mean field width of the grid cell’s response. (3) The widest field of the response must be smaller than 5w. (4) At least 30% of the bins must be assigned to either in-field or out-of-field periods. (5) The ratio between in-field and out-of-field mean ΔF/F needs to be ≥ 2^68,92,93^. (6) The cell must not be identified as a cue cell on that day. Common cells were identified as true grid cells if they were identified as grid cells on more than half of imaging days (≥ 6). All cells that were not identified as grid cells were considered non-grid cells. Non-grid cells are distinct from unclassified cells (see “Unclassified Cell Identification”)

##### Speed Score/Speed Cell Classification

Speed score was calculated as previously described^37,38^. First, spatial position for the full session was smoothed using a 0.8s gaussian filter. Velocity was calculated using the smoothed positions as the difference in position divided by the difference in time. Time points with velocity less than zero or greater than 100 cm/s were ignored in the rest of the calculation. A histogram of speed was then used to determine the largest 10 cm/s speed range (0-10 to 90-100) that the mouse occupied for at least 30 s, and the mean velocity of time points with speed in the range was used as a new upper speed limit. A new lower limit of 6 cm/s was also used. If the new upper limit was lower than 6 cm/s, the upper limit was adjusted using the averaged velocity of the next 10 cm/s speed bin. Time points with velocity outside these new limits were ignored in the rest of the calculation. The temporal ΔF/F trace was smoothed with a 0.4s gaussian filter. Raw speed score is defined as the Pearson correlation between velocity and ΔF/F for time points with velocity in the final range. For calculating positive and negative speed score components, classification was performed daily, so a given cell could contribute to the positive speed score component on one day and the negative speed score component on another. Absolute speed score is the absolute value of the raw speed score.

1000 shuffled speed scores were generated for each cell using the same calculation but the temporal ΔF/F trace was rotated as described above (see Spatial Field Identification). Shuffled speed scores from all cells across all FOVs and imaging days were pooled to generate an overall speed score distribution. A cell was identified as a speed cell if its speed score on the day was above the 99th percentile (positive speed cell) or below the 1^st^ percentile (negative speed cell) of the shuffle cue score distribution. Speed cell identification was performed on each day. Common cells were identified as true speed cells if they were identified as speed cells of either type (positive or negative) on more than half of imaging days (≥ 6).

##### Unclassified Cell Identification

Unclassified cells were all common cells that were not identified as cue, grid, or speed cells. Note that cells could not be classified as both a cue cell and a grid cell, but cells could be classified as a speed cell and either a cue cell or a grid cell. Therefore, the combined number of cue, grid, speed, and unclassified cells could be greater than the total number of common cells.

##### Within-Day Consistency

The within-day consistency for a given cell in a particular imaging session was calculated as previously described^98^. The spatially binned mean ΔF/F for each lap along the track was correlated with that of every other lap. The average of these correlations is the within-day consistency value for a given cell.

##### Cross-Day Consistency

The activity matrix correlation (cross-day consistency) for a given set of cells in a particular imaging session was calculated as previously described^49^. The spatially binned mean ΔF/F for each cell was averaged across all laps along the track, generating a 1D array. The calculated array for each cell was concatenated to generate a single matrix (size = total number of cells by number of spatial bins) such that the activity of a given tracked cell was in the same row of the activity matrix for all days. The day-to-day activity matrix correlation was calculated on a per cell basis as the 1D Pearson correlation between the generated 1D arrays for a given cell on two consecutive days.

##### Inter-Mouse Consistency

To calculate inter-mouse consistency, a spatial map for each mouse on a given day or set of days was calculated. The spatially binned mean ΔF/F for all common cells from both FOV from a given mouse were averaged to generate a spatial map for each day. The daily maps (NE days 7-10) were averaged together to generate the late-learning spatial map for each mouse. Inter-mouse consistency between a pair of mice was defined as the correlation between the daily or late-learning maps of those mice. Within-group and between-group comparisons were those between two mice of the same or different sex/genotype, respectively, as described in the text. To calculate the difference between WT and PS19 mice for the sex versus molecular cell type heatmap and to calculate the values used for the general linear models predicting behavior, the correlations of each mouse with every other mouse were averaged to generate a single value per mouse per day or per set of days.

##### Position Decoding

Position decoding was performed by separating the imaging data into template data (odd laps) and testing data (even laps) as described previously^49,98^. For this analysis, the raw ΔF/F (not only significant transients) was used because performing the decoding analysis with the significant transient ΔF/F resulted in high decoding accuracy at the ends of the track. Because we were interested in decoding as a function of track position, we used raw ΔF/F rather than excluding the edge of the track. The spatially binned (5 cm bins) mean ΔF/F for the template data laps were averaged across laps and concatenated for all cells to generate a template matrix (size = total number of cells by number of spatial bins). The ΔF/F of all cells at each time were then correlated to each spatial bin of the averaged template data matrix. The decoded position for a given time point was the spatial position of the template bin that gave the highest correlation. Decoding accuracy was the percentage of correctly decoded track positions, which was within 2 spatial bins from the spatial bin of the real position. The first and last spatial bin of the track were considered adjacent for this calculation. The decoding was performed with 100 random samples of 30 cells for each FOV. All cells or all cells identified as grid cells or non-grid cells on a given day, were used for this analysis due to the small number of common cells in some FOV. 30 cells were chosen for the analysis to maximize the number of FOV with sufficient cells (>∼90% of FOV had 30 cells for each cell category. All Cells: 412/412 FOV had 30 cells, Grid Cells: 366/412 FOV had 30 cells, Non-Grid Cells: 403/412 FOV had 30 cells). This analysis could not be separately conducted on stellate or pyramidal cells because many FOV had too few cells in a particular cell category.

##### Spatial Activity Cue Anchoring and Decoding Cue Anchoring

The spatially binned mean ΔF/F for all common cells from a given FOV were averaged to generate a spatial map for each day (spatial activity level). This calculation was performed per FOV to account for the noise in this measurement. Spatial position decoding was defined as the decoding accuracy as described above averaged for test bins at each spatial position. This resulted in a 1D array representing position decoding as a function of track position (spatial decoding).

For spatial activity level and spatial decoding, in-cue regions were defined as the average of all spatial bins within an expanded cue region (the cue itself and the surrounding region expanded by half of the cue width on both sides). Out-cue regions were defined as the average of all spatial bins not part of the in-cue region. The full track decoding was the average spatial decoding across the full track. Spatial activity cue anchoring and decoding cue anchoring were defined as the in-cue/out-cue (I/O) ratio, or the in-cue value divided by the out-cue value.

##### General Linear Model

###### Model Inputs

As described in the text, the predictor variables included behavioral variables (mean velocity), general activity variables (activity level, spatial selectivity, raw speed score, and absolute speed score), map consistency variables (within-day consistency, cross-day consistency, and inter-mouse consistency), and global representation variables (spatial activity cue anchoring and decoding cue anchoring). For activity variables calculated by FOV (spatial activity cue anchoring and decoding cue anchoring), the average activity value of the two FOVs for the given cell type was used as the input for a given day. For activity variables calculated by mouse (inter-mouse consistency, as described above), the daily activity value was used directly. For activity variables calculated by cell (all others), the average activity of all cells for a given cell type from both imaging FOV was used as the input for a given day. As described above, decoding cue anchoring was calculated using all cells rather than common cells and could not be calculated for stellate and pyramidal cells. Because all other activity variables were calculated using common cells, the model types will be referred to as all-common, grid, non-grid, stellate, and pyramidal even though decoding cue anchoring was calculated using all cells. Decoding cue anchoring was not used as a predictor variable when comparing the stellate and pyramidal models to the all-common-cell model. Sex and genotype were included as confounding variables. Mouse identity and session number were not included in the model. All predictor variables were standardized using z-score normalization.

All modeling was performed using a general linear model (GLM)^101^ using the MATLAB stepwiseglm and fitglm functions. Because success on each lap was binary and all mice experienced 19 laps that were used for behavioral analysis each NE day, the GLM always used a binomial distribution with a binomial size of 19.

###### Variable Selection

Variable selection was performed by iteratively removing predictor variables from a full model using the MATLAB stepwiseglm function to eliminate predictor variables that did not provide significant contributions to the model. This was performed only for the all-common-cell data. The initial model formula included all linear terms with no interactions, so all predictor and confounding variables were included. The input for the upper bound of the model was also all linear terms with no interactions to prevent nonlinear or interaction terms being added. The input for the lower bound of the model included just the confounding variable terms to prevent these variables from being removed. The maximum number of steps was set so that at least one predictor term would always remain.

This procedure was performed using leave-one-out cross-validation (LOOCV)^186^ across mice. Each iteration, 20 mice were used for training and the one left-out mouse was then used for testing. A total of 21 iterations were performed so that each mouse served as the left-out mouse once, and 21 sets of variables were obtained from the iterative removal process. A final variable set was selected using only those variables that were present in all 21 post-removal variable sets. The same procedure was repeated but excluding sex and genotype to determine whether the confounding variables affected variable selection.

To test the validity of this iterative removal process, the same procedure was performed when the success rate was shuffled across mice and days to remove the correlation between activity and behavior. For both the real data and the shuffled data, the predictions for the test mouse from all 21 iterations were combined to generate a final combined prediction. An R^2^ goodness of fit (variance explained) was calculated between the predicted behavior and the real or shuffle behavior. A total of 1,000 shuffles of success rate were performed to generate a distribution of variance explained in the absence of any connection between the predictors and success rate. The p-value of the iterative removal method was defined as (R^2^_greater_ + R^2^ /2)/(nShuffle + 1), where R^2^ is the number of shuffle R^2^ values greater than the real data R^2^ value, R^2^ is the number of shuffle R^2^ values equal to the real data R^2^ value, and nShuffle is the number of shuffles performed^102^.

###### Final Model Training and Evaluation

To evaluate the selection of the final variables determined above and the predictive ability of these variables, a final model was trained on only the final variable data using the MATLAB fitglm function. LOOCV was again performed for this analysis and those described below in the same manner as above. The final model coefficients were the average coefficients of the 21 LOOCV iteration models.

To evaluate the final variable selection, this final model was compared to sub-models trained with each of the final variables removed one at a time and to expansion-models with each of the variables not in the final model added one at a time^187^. The training R^2^ goodness of fit is calculated for the final model, each sub-model, and each expansion-model by averaging the training predictions of each mouse across the 20 LOOCV iteration that it was a training mouse, combining these averaged training predictions across mice, and comparing to the true behavior.

To directly evaluate the final model fit, a testing R^2^ goodness of fit was calculated by combining the predictions for the test mouse from all 21 iterations and comparing to the true behavior. To evaluate the ability of the model to predict learning behavior, the prediction time courses of individual mice or groups were compared to their true behavior. Individual mouse prediction time courses were the test prediction on the LOOCV iteration when that mouse was the test mouse. The prediction time course for each group was calculated as the average of the test predictions across all mice in that group. The true behavior time course for each group was the average success rate across all mice in that group.

###### Dominance Analysis

To evaluate the relative contribution of each of the final variables to the final model prediction, dominance analysis was performed^103^. We determined the effect of removing each variable from the full model and from all possible sub-models to determine the average contribution of each variable^103^. LOOCV is again used, with each mouse being used as the test mouse once. For each LOOCV iteration, the full model and every possible sub-model are trained using the MATLAB fitglm function. In other words, a GLM is trained using every possible combination of the variables (activity level, spatial selectivity, absolute speed score, within-day consistency, decoding cue anchoring, and spatial activity cue anchoring, sex, and genotype).

To evaluate the contribution of a given variable, the training R^2^ goodness of fit of all sub-models with this variable are compared to the smaller sub-models where this variable is removed. For example, if a full model has variables A, B, and C, the contribution of the variable A is made by comparing the model ABC to BC, AB to B, AC to C, and A to the coefficient 1. A tier is defined as all sub-models with a specific number of variables (i.e. models with only two variables). The comparisons made for all sub-models in a tier are averaged together to generate a final tier R^2^ value. For example, tier 2 of the 3 variable model has the possible sub-models AB and AC when evaluating the contribution of the variable A. Therefore, the tier 2 R^2^ value for the variable A would be the average of the comparisons for AB to B and AC to C.

A full model with N variables will have N tiers of R^2^ values for each variable. The final variance explained for a variable is the average of the R^2^ values for all N tiers. The LOOCV iterations are combined by averaging the final variance explained for each variable across iterations. The sum of the variance explained across all variables is roughly equal to variance explained by the full model^103^, and can therefore be thought of as the relative contribution to the final variance explained.

###### Model Input Subsampling

Activity data was subsampled to match the number of cells that contributed to the activity in each mouse across cell types (all-common vs. grid vs. non-grid or all-common vs. stellate vs. pyramidal). The minimum number of cells across cell types was determined for each mouse. For each subsample iteration (N = 200 iterations), that number of cells minus 1 were randomly selected across the two FOV for that mouse and the activity parameters of only those cells were calculated. Because decoding cue anchoring already used 30 cells per FOV, the original values of this variable were used. The above procedure (“Final Model Training and Evaluation”) was used to generate an R^2^ variance explained for the test predictions of each subsample iteration for each cell type. Power analysis to determine the power of the difference between groups (the probability of obtaining a significant result if a true difference exists) if the number of subsamples was varied was performed using the MATLAB sampsizepwr function assuming a two-sample pooled t-test. No additional exclusions were made comparing the all-common, grid, and non-grid models. As mentioned above, decoding cue anchoring was not used for the comparison of the all-common, stellate, and pyramidal models. In addition, because several mice had very few stellate or pyramidal cells, mice with fewer than 3 cells were excluded from the analysis.

###### Genotype Classification

To classify mouse genotype based on the behavior predictions of MEC neuronal activity, the above procedure (“Final Model Training and Evaluation”) was used to generate test predictions for each cell type. Because these results were not compared across cell types, no subsampling was performed. Instead all common cells or all grid, non-grid, stellate, or pyramidal cells among the common cell population were used (except for decoding cue anchoring, which used all cells, all grid cells, or all non-grid cells). As above, decoding cue anchoring was not used for the stellate or pyramidal cell models. In addition, sex and genotype were not included to avoid biasing the classification of genotype. Mice with fewer than 3 cells of a given cell type were excluded from the analysis of that cell type.

The average test prediction across late learning (days 7-10) of each mouse was classified as WT or PS19 based on whether it was closer to the average true behavior of WT or PS19 mice of that sex. In the rare event of a tie, classification was considered half correct (including a half contribution to the classification accuracy below). Each mouse was excluded from the average true behavior calculations for its own test prediction, except for in the pyramidal cell model, which used all mice to calculate the average true behavior due to the smaller number of mice included in the calculation. The classification accuracy of a given model for a given sex was the percentage of mice of both genotypes combined that were correctly classified. An identical shuffle procedure was used to that described above (200 shuffles, see “Variable Selection”) to generate a shuffle distribution of genotype classification accuracy. The percentile of the classification of real predictions relative to the shuffles was reported.

#### Activity Heatmaps

The 2-by-2 heatmaps to visualize the interaction between molecular cell identity (stellate vs. pyramidal) and by sex for all activity parameters were generated by taking the difference between the average WT and PS19 values (or PS19 and WT when high activity values are considered a deficit, as noted in the figures) and the significance corresponds to an unpaired t-test. The 1-by-3 heatmaps for behavior are generated using the same method. The 2-by-2 heatmap for pTau overlap represents the stellate or pyramidal cell overlap with pTau. The 2-by-2 heatmap for pTau intensity represents the mean intensity of each sex/cell type group.

The averaged tau accumulation and activity heatmaps were generated using all tau accumulation or activity data heatmaps (those in **Figs. 6A, B**). To standardize the scale of the heatmaps, each tau accumulation or activity heatmap was normalized by the maximum of the absolute value of the heatmap values, so the values of all the heatmaps ranged between -1 and 1. These standardized heatmaps were averaged to generate the final average heatmaps.

#### General data analysis and statistics

Image processing was performed using previously published MATLAB (MathWorks, version R2015aSP1) codes as cited above and Suite2p (v0.11.1)^183^. Data analysis was performed using ImageJ (Fiji, version 1.53q), MATLAB (MathWorks, versions R2015aSP1 and 2020a), and GraphPad Prism (version 9.3.1). Linear correlations and the corresponding r and p values were calculated using Pearson’s linear correlation. Significance values for comparing between group means were calculated using unpaired Student’s t test, paired Student’s t test, or ANOVA tests (two-way ordinary or repeated measures ANOVA; sphericity is assumed for all tests), or Linear Mixed-Effects model (REML, random intercepts for subjects, missing values were excluded, categorical predictors were effects-coded, significance was tested via ANOVA with Satterthwaite’s method) as noted. Where noted, multiple pairwise comparisons were performed following ANOVA tests using unpaired Student’s t test. P values were adjusted for multiple comparisons using the Bonferroni–Holm method as noted. For the survival curve analysis, groups were compared using a log-rank (Mantel-Cox) test. For fitting success rate to one-phase association exponential functions, an extra sum-of-squares F test was used to select between model fits as described above. For this test, a p-value of 1 indicates the simpler model fit the data better than the more complex model, and so the F test was not performed. Significance values for averaged activity heatmap used one-sample Student’s t-test, comparing the normalized activity values within each square to 0. Two-tailed tests were used for all analyses. P values less than 0.05 were considered significant (* < 0.05, ** <0.01, *** <0.001). All line plots show mean and standard error. In all violin plots, horizontal line represents mean, white circle represents median, and whiskers represent interquartile range. Violins represent the kernel density estimation of the data. Detailed statistical information for all figures can be found in **Table S1**.

